# Spatiotemporal prediction of wildfire extremes with Bayesian finite sample maxima

**DOI:** 10.1101/384115

**Authors:** Maxwell B. Joseph, Matthew W. Rossi, Nathan P. Mietkiewicz, Adam L. Mahood, Megan E. Cattau, Lise Ann St. Denis, R. Chelsea Nagy, Virginia Iglesias, John T. Abatzoglou, Jennifer K. Balch

**Affiliations:** Earth Lab, University of Colorado Boulder; Department of Geography, University of Idaho

**Keywords:** fire, wildfire, Bayesian, spatiotemporal, extremes, climate

## Abstract

Wildfires are becoming more frequent in parts of the globe, but predicting where and when wildfires occur remains difficult. To predict wildfire extremes across the contiguous United States, we integrate a 30 year wildfire record with meteorological and housing data in spatiotemporal Bayesian statistical models with spatially varying nonlinear effects. We compared different distributions for the number and sizes of large fires to generate a posterior predictive distribution based on finite sample maxima for extreme events (the largest fires over bounded spatiotemporal domains). A zero-inflated negative binomial model for fire counts and a lognormal model for burned areas provided the best performance. This model attains 99% interval coverage for the number of fires and 93% coverage for fire sizes over a six year withheld data set. Dryness and air temperature strongly predict extreme wildfire probabilities. Housing density has a hump-shaped relationship with fire occurrence, with more fires occurring at intermediate housing densities. Statistically, these drivers affect the chance of an extreme wildfire in two ways: by altering fire size distributions, and by altering fire frequency, which influences sampling from the tails of fire size distributions. We conclude that recent extremes should not be surprising, and that the contiguous United States may be on the verge of even larger wildfire extremes.

## Introduction

Wildfire frequency and burned area has increased over the past couple decades in the United States (Dennison et al. 2014; Westerling 2016), and elsewhere (Krawchuk et al. 2009; Pechony and Shindell 2010). In addition to the ecological and smoke impacts associated with increased burned area, there has been an increasing interest in extreme wildfires (Williams 2013) given their impact on human lives and infrastructure (Kochi et al. 2010; Diaz 2012). While case studies of particular extremes provide insight into what caused past events (Peterson et al. 2015; Nauslar, Abatzoglou, and Marsh 2018), predictions of future extremes at a national level could inform disaster related resource allocation. The term “extreme” has multiple meanings with respect to wildfires (Tedim et al. 2018), and in this paper we take consider an extreme wildfire to be a fire with the largest burned area over a bounded spatiotemporal domain, i.e., the block maximum within a spatial region and a temporal interval (Coles et al. 2001). For example, the block maxima for widlfires across the contiguous U.S. can be defined on a yearly basis (Figure 1).

**Figure 1.**
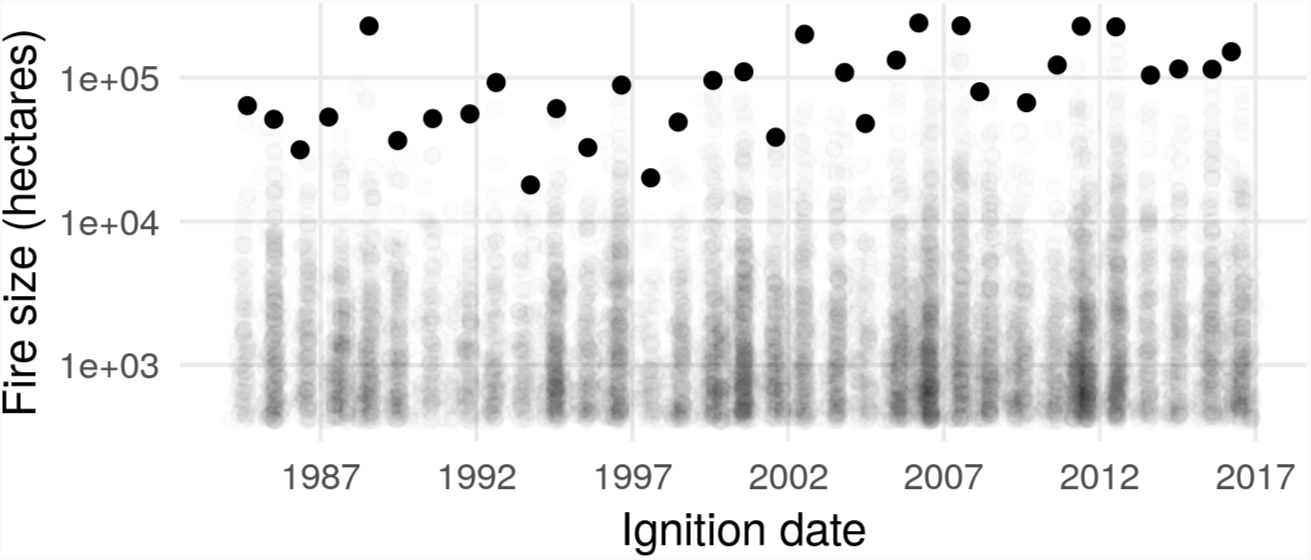
Sizes of wildfires over 405 hectares in the contiguous United States, from the Monitoring Trends in Burn Severity multiagency program. Each point represents a fire event, and the largest fires for each year (the block maxima) are shown as solid black points.

Factors driving wildfire extremes vary in space and time (Barbero et al. 2014), but it is unclear how best to account for this in a predictive model. Previous efforts have used year or region-specific models, aggregating over space or time (Bermudez et al. 2009), temporally or spatially explicit models (Mendes et al. 2010), and spatial models with year as a covariate (Díaz-Avalos, Juan, and Serra-Saurina 2016). Recently, rich spatiotemporal models have been described with linear, spatially constant covariate effects (Serra, Saez, Juan, et al. 2014; Serra, Saez, Mateu, et al. 2014). However, linear, spatially constant effects are suboptimal over large spatial domains with nonlinear drivers (Fosberg 1978, Goodrick (2002), Preisler et al. (2004); Preisler and Westerling 2007; Balshi et al. 2009; Krawchuk et al. 2009; Pechony and Shindell 2009; Vilar et al. 2010; Woolford et al. 2011; Woolford et al. 2014). For example, global wildfire probability shows a hump-shaped relationship with temperature and moisture (Moritz et al. 2012). Interactions among drivers also impose nonlinearity, e.g., in hot and dry climates fires are fuel limited (McLaughlin and Bowers 1982), but in cold and wet climates fires are energy limited (Krawchuk and Moritz 2011).

Prediction is also complicated by uncertainty in which distribution(s) to use to assign probabilities to extreme events. The generalized Pareto distribution (GPD) has frequently been used (Bermudez et al. 2009; Jiang and Zhuang 2011), but the GPD requires a threshold to delineate extreme events (Davison and Smith 1990, Coles (2014)). The utility and validity of a threshold for extremes in a heterogeneous region is debatable (Tedim et al. 2018). Recently proposed metastatistical extreme value (MEV) approaches do not require such a threshold, and are based on the statistical distribution of finite sample maxima, i.e., the probability distribution of the maximum value for a finite number of events (Marani and Ignaccolo 2015; Zorzetto, Botter, and Marani 2016). In the MEV framework, the occurrence and size of future events, and the parameters of their distributions are treated as random variables which together imply a distribution for extremes. This approach has roots in compound distributions (Dubey 1970; Wiitala 1999), doubly stochastic processes (Cox and Isham 1980), superstatistics (Beck and Cohen 2003), and the Bayesian posterior predictive distribution (Gelman et al. 2013). The link to Bayesian inference is particularly useful, as it provides an easy way to propagate uncertainty forward to to predictions of extremes (Coles, Pericchi, and Sisson 2003).

Here, we extend the finite sample maximum approach to account for non-linear, spatially varying covariate effects with the goal of predicting extreme wildfire events from a statistical perspective across the contiguous United States. Specifically, we aim to predict occurrence (where and when), and magnitude (burned area) of large wildfires at a monthly time scale and regional spatial scale across the contiguous United States.

## Methods

### Data description

We acquired wildfire event data for the contiguous United States from the Monitoring Trends in Burn Severity (MTBS, www.mtbs.gov) program (Eidenshink et al. 2007), which includes spatiotemporal information on the occurrence of wildfires in the United States from 1984 to 2016. The MTBS data contain fires greater than 1000 acres (≈ 405 hectares) in the western U.S. and greater than 500 acres (≈ 202 hectares) in the eastern U.S. For consistency across the U.S., we discarded all records in the MTBS data less than 1000 acres, retaining 10,736 fire events (Figure 2A). Each event in the MTBS data has a discovery date, spatial point location, and final size.

**Figure 2.**
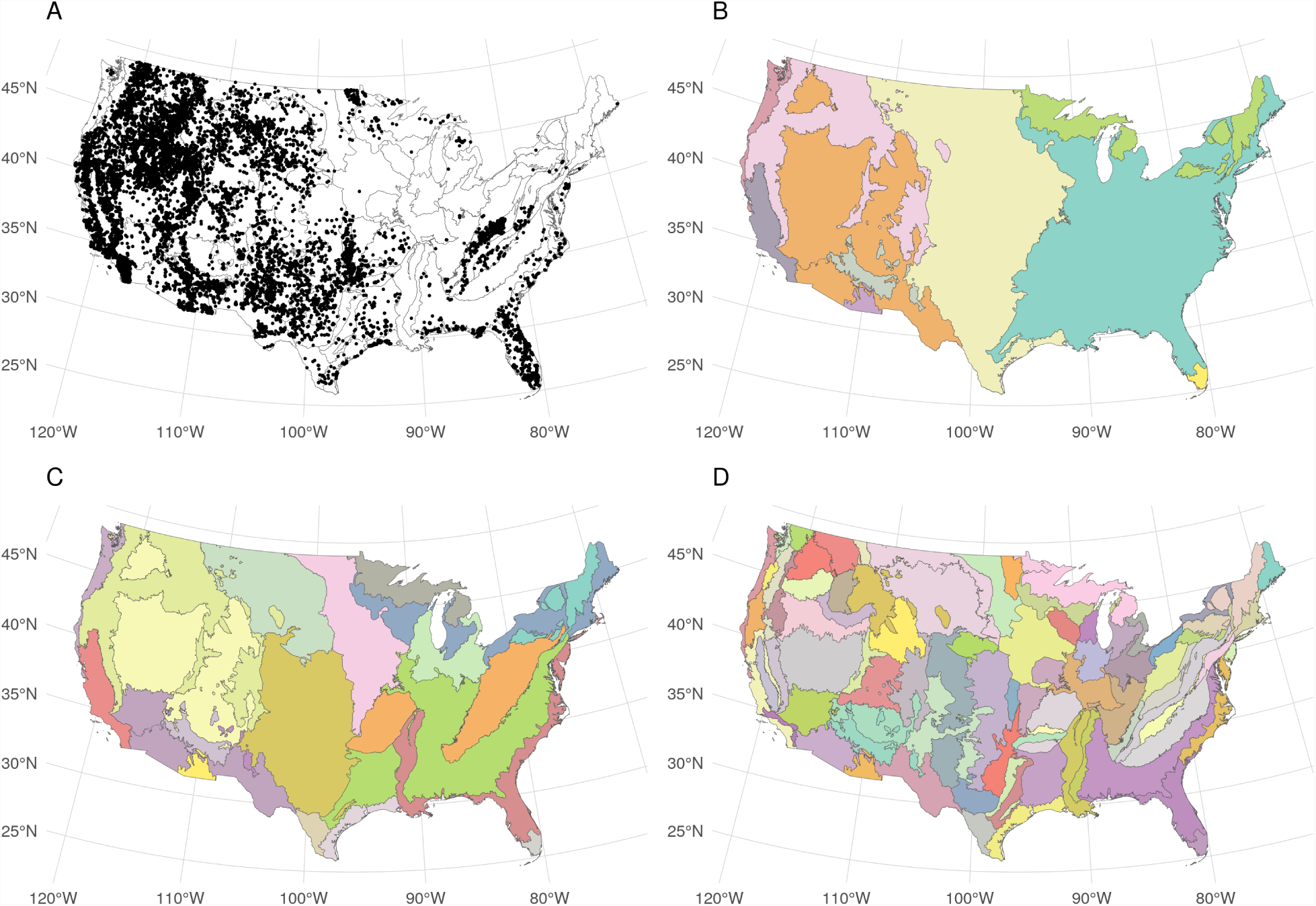
A. Large wildfire ignition locations are shown as points across the study region. Colors in panels B, C, and D show level 1, 2, and 3 ecoregions respectively.

To explain fire size and occurrence, we used a combination of meteorological variables including humidity, air temperature, precipitation, and wind speed. These variables were selected on the basis of previous work, and also with an aim to drive a predictive model with interpretable meteorological quantities. Meteorological layers were acquired from the gridMET data (Abatzoglou 2013) that blends monthly high-spatial resolution (∼4-km) climate data from the Parameter-elevation Relationships on Independent Slopes Model (Daly et al. 2008) with high-temporal resolution (hourly) data from the National Land Data Assimilation System (NLDAS2) using climatologically aided interpolation. The resultant products are a suite of surface meteorological variables summarized at the daily time step and at a 4-km pixel resolution. Daily total precipitation, minimum relative humidity, mean wind speed, and maximum air temperature were averaged at a monthly time step for each of 84 Environmental Protection Agency level 3 (L3) ecoregions for each month from 1984 to 2016 (Omernik 1987; Omernik and Griffith 2014). We also computed cumulative monthly precipitation over the previous 12 months for each ecoregion-month combination. We chose to segment the U.S. with level 3 ecoregions as a compromise between the more numerous (computationally demanding) level 4 ecoregions, and the coarser level 2 ecoregions.

We used publicly available housing density estimates that were generated based on the U.S. 2000 decennial census as explanatory variables that may relate to human ignition pressure (Radeloff et al. 2010). These are provided at decadal time steps, and spatially at the level of census partial block groups. To generate approximate measures of housing density at monthly time intervals, we used a simple linear interpolation over time for each block group, then aggregated spatially across block groups to compute mean housing density for each ecoregion in each month.

### Model development

We built two types of models: one describing the occurrence of fires within each L3 ecoregion over time (i.e., the total number of fires occurring in each ecoregion for each month from 1984 - 2016), and another describing the size of each wildfire in each ecoregion and month. For occurrence models, the response variable was a count (number of fires), and for burned area models, the response was a continuous positive quantity (size of each fire event). We used the period from 1984 to 2009 for training, witholding the period from 2010 to 2016 to evaluate predictive performance.

#### Fire occurrence

We constructed four models for fire occurrence and compared their predictive performance based on test-set log likelihood and posterior predictive checks for the proportion of zeros, maximum count, and total count. The models differed in the distributions used in the likelihood, representing counts as a Poisson, negative binomial, zero-inflated Poisson, or zeroinflated negative binomial random variable. The Poisson distribution is a common choice for counts, and the negative binomial distribution provides an alternative that can account for overdispersion. The zero-inflated versions of these distributions include a component to represent extra zeros, which might be expected to work well if there are independent processes that determine whether nonzero counts are possible (Lambert 1992).

For spatial units (ecoregions) *s* = 1, *…, S* and time steps (months) *t* = 1, *…, T*, each model defines a probability mass function for *n_s, t_*: the number of fires over 405 hectares in ecoregion *s* and time step *t*. For each of the four count distributions under consideration, location parameters *µ*_*s, t*_ and (for zero-inflated models) structural zero inflation parameters *π*_*s, t*_ were allowed to vary in space and time. We used a log link function to ensure that *µ*_*s, t*_ *>* 0, and a logit link function to ensure that *π*_*s, t*_ ∈ (0, 1). Concatenating over spatial and temporal units, so that ***µ*** = (*µ*_*s*=1, *t*=1_, *µ*_*s*=2, *t*=1, *…,*_ *µ*_*s*=*S, t*=1_, *µ*_*s*=*S, t*=2, *…,*_ *µ*_*s*=*S, t*=*T*_), and similarly for ***π***, we modeled location and (when applicable) zero inflation parameters as:

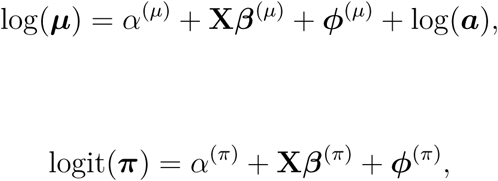

where α ^(*µ*)^ andα ^(*π*)^ are scalar intercept parameters, **X** is a known (*S × T*) *× p* design matrix, where *p* is the number of input features, *β* ^(*µ*)^ and *β*^(*π*)^ are column vector parameters of length *p, ϕ*^(*µ*)^ and *ϕ*^(*π*)^ are column vector parameters of length *S × T* containing spatiotem-poral adjustments, and ***a*** is a known offset vector of areas for spatial unit *s* = 1, 2, *…, S*, repeated *T* times.

#### Burned area

We developed five candidate models for fire size, each of which specified a different distribution for the size (burned area) of individual fire events (Reed and McKelvey 2002; Hernandez et al. 2015), including the generalized Pareto (Hosking and Wallis 1987), tapered Pareto (Schoenberg, Peng, and Woods 2003), lognormal, gamma, and Weibull distributions. We evaluated each model in terms of test set log likelihood, and posterior predictive checks for fire size extremes. We defined the response *y*_*i*_ as the number of hectares burned over 405 for the *i*^*th*^ fire event, which occurred in spatial unit *s*_*i*_ and time step *t*_*i*_.

Because each burned area distribution has a different parameterization, we included covariate effects in a distribution-specific way. For the generalized Pareto distribution (GPD), we assumed a positive shape parameter, leading to a Lomax distribution for exceedances (Bermudez et al. 2009). The GPD and Lomax shape parameters are related by *κ*^(*GPD*)^ = 1*/κ*^(*L*)^, and the GPD scale parameter is related to the Lomax scale and shape parameters by *σ*^(*GPD*)^ = *σ*^(*L*)^*/κ*^(*L*)^. We introduced covariate dependence via the Lomax scale parameter using a log link. For event *i*, 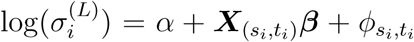, where *α* is an intercept parameter, ***β*** is a length *p* vector of coefficients, 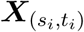 is a row vector from **X**, and 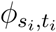 is a spatiotemporal adjustment for *s*_*i*_ and *t*_*i*_. For the tapered Pareto model, we modeled the shape parameter as 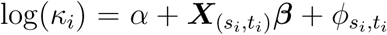. The lognormal model included covariate dependence via the location parameter: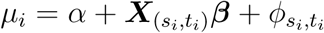. The gamma model used a log link for the expected value: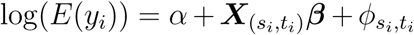. Last, we modeled the Weibull scale parameter as 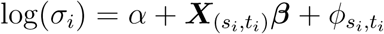. More detail on the parameterization of each burned area distribution is provided in the Appendices.

#### Accounting for nonlinear forcing

The design matrix **X** was constructed to allow for spatially varying nonlinear effects of housing density and meteorological drivers. We used B-splines to account for nonlinearity (Figure 3) and allowed the coefficients for each basis vector to vary spatially (Wood 2017). First, we constructed univariate B-splines for log housing density, wind speed, same month precipitation, previous 12 month precipitation, air temperature, and humidity, with five degrees of freedom (including an intercept) for each variable. This step generated 30 basis vectors (five for each of six variables).

**Figure 3.**
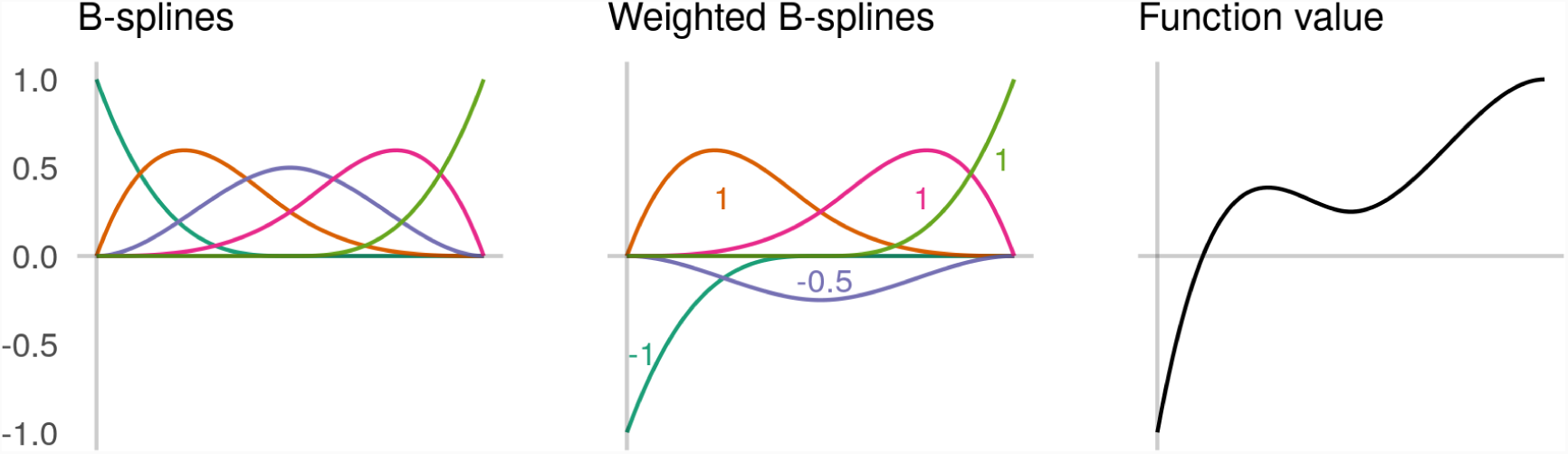
Conceptual figure to illustrate the use of B-splines to construct nonlinear functions. In the left panel, five B-spline vectors are shown, which map values of an input variable (on the x-axis) to a value on the y-axis. The middle panel shows the same B-spline vectors, but weighted (multiplied) by real numbers, with the weights illustrated as annotations. These weighted B-spline vectors are summed to produce the values of a nonlinear function (right panel).

To allow for spatial variation in these nonlinear effects, we added interaction effects between each of the basis vectors and ecoregions (Brezger and Lang 2006; Kneib, Hothorn, and Tutz 2009). The hierarchical nesting of ecoregion designations (Figure 2B-D) lends itself to such interactions. Conceptually, coefficients in a level 3 ecoregion may be related to coefficients in the level 2 ecoregion containing the level 3 region, the level 1 ecoregion containing the level 2 region, and a global effect. The coefficient associated with a basis vector for any level 3 ecoregion is treated as a sum of a global effect, a level 1 ecoregion adjustment, a level 2 ecoregion adjustment, and a level 3 ecogregion adjustment. Thus, for every univariate basis vector, we included interaction effects with ecoregion at each of the three ecoregion levels. This allows borrowing of information across space (level 3 ecoregions in a level 2 ecoregion are often adjacent), and for regions that are ecologically similar. We also included adjustments on the global intercept for each level 1, 2, and 3 ecoregion to account for spatial variation that is unrelated to climate or housing density. This specification induces sparsity in **X** that we exploit to increase the efficiency of computing ***µ*** and ***π***. In total, **X** has *p* = 3,472 columns, with 97% zero entries.

### Prior specification

To avoid overfitting, we used a regularized horseshoe prior on the coefficients associated with the spatially varying nonlinear effects described above (Piironen, Vehtari, and others 2017). This prior places high probability close to zero, while retaining heavy enough tails that nonzero coefficients are not shrunk too strongly toward zero. This is consistent with our prior expectation that most of the coefficients associated with the columns in **X** were close to zero. For the zero inflated count models, we used a multivariate horseshoe to allow information sharing between the zero inflated and distribution specific location parameters (Peltola et al. 2014). For the remaining count models and all burned area models, this was a univariate horseshoe prior. Spatiotemporal random effects were constructed using a temporally autoregressive, spatially intrinsically autoregressive formulation (Besag and Kooperberg 1995; Banerjee, Carlin, and Gelfand 2014). Details of these priors and the resulting joint distributions are provided in the Appendices.

### Posterior predictive inference for finite sample maxima

We used the posterior predictive distribution to check each model and make inference on extremes. The posterior predictive distribution provides a distribution for replications of observed data (*y*^rep^), and predictions of future data (Gelman et al. 2013). Conceptually, for a good model, *y*^rep^ should be similar to observed training data *y*, and future predictions should be similar to future data. Distributions over both quantities can be obtained by conditioning on *y* and marginalizing over model parameters *θ*, e.g., [*y*^rep^*|y*] =∫ [*y*^rep^*|θ*][*θ|y*]*dθ*.

Posterior predictive distributions facilitate model checks that compare predicted and observed test statistics (Gelman, Meng, and Stern 1996). To evaluate whether models captured tail behavior, we compared empirical maxima (*T* (*y*) = max(*y*)) to the predicted distribution of maxima *T* (*y*^rep^). We also include predictive checks for the proportion of zero counts, and totals for count and burned area models. Posterior predictive inference for finite sample maxima is similar in spirit to the MEV approach. Both obtain a distribution over maxima by marginalizing over unknowns including the number of events, size of each event, and parameters of their distributions (Marani and Ignaccolo 2015). However, a Bayesian approach explicitly conditions on the observed data to obtain a posterior distribution of parameters.

Seeing this connection is useful in the context of including priors and propagating uncertainty to derived parameters. For any ecoregion *s* and timestep *t*, if we define a particular maximum fire size conditional on a fire having occurred as *z*_*s, t*_, and let *Z*_*s, t*_ represent the random variable of maximum fire size, then the cumulative distribution function (CDF) for *z*_*s, t*_ is given by 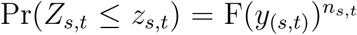, where F(*y*_(*s, t*)_) is the CDF of fire size, and *n*_*s, t*_ is the number of wildfire events. Thus, Pr(*Z*_*s, t*_ *≤ z*_*s, t*_) is the distribution function for the finite sample maximum. The CDF for *z*_*s, t*_ can be inverted to produce a quantile function that permits computation of prediction intervals for maximum fire sizes, conditional on fires having occurred. Given a collection of posterior draws from a burned area model that parameterize F(*y*_(*s, t*)_), and a collection of posterior draws of 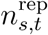 from the posterior predictive distribution of a wildfire count model, a posterior distribution for the CDF or quantile function of maximum fire size can be generated which combines the two models to facilitate inference on the distribution of extremes.

### Parameter estimation

We used a combination of variational approximations and Hamiltonian Monte Carlo methods to sample from the posterior distributions of count and burned area models. A variational approximation (Kucukelbir et al. 2015) was used for count models to quickly identify a preferred model. The best performing count model and all burned area models were fit using the No-U-Turn Sampler (Hoffman and Gelman 2014). Models were fit in the Stan probabilistic programming language using the `rstan` package (Carpenter et al. 2016; Stan Development Team 2018). We ran four chains for 1000 iterations each, discarding the first 500 iterations as warmup. Convergence was assessed using visual inspection of trace plots, with potential scale reduction statistic values 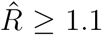 as an indicator convergence failure (Brooks and Gelman 1998).

### Implementation

All data processing, model fitting, and visualization were implemented with open source software, primarily in the R programming language (R Core Team 2017), and wrapped in a reproducible workflow via GNU Make and Docker (Stallman, McGrath, and Smith 2004; Boettiger 2015). Data cleaning and transformation required the R packages assertthat (Wickham 2017a), lubridate (Grolemund and Wickham 2011), Matrix (Bates and Maechler 2018), pbapply (Solymos and Zawadzki 2018), splines (R Core Team 2018), tidyverse (Wickham 2017b), and zoo (Zeileis and Grothendieck 2005). Spatial data were processed with raster (Hijmans 2017), rgdal (Bivand, Keitt, and Rowlingson 2018), sf (Pebesma 2018), and spdep (Bivand and Piras 2015). Finally, we used cowplot (Wilke 2017), ggrepel (Slowikowski 2018), ggthemes (Arnold 2018), patchwork (Pedersen 2017), and RColorBrewer (Neuwirth 2014) for visualization. The manuscript was written in R Markdown (Allaire et al. 2018). Analyses were run on an Amazon Web Services m5.2xlarge EC2 instance with four physical cores and 32 GB of RAM, and the whole workflow requires ≈ 72 hours. All code to reproduce the analysis is available on GitHub at https://github.com/mbjoseph/wildfire-extremes (Joseph 2018).

## Results

### Wildfire occurrence

The zero-inflated negative binomial distribution performed best on the held-out test set (Table 1), and was able to recover the proportion of zeros, count maxima, and count totals in posterior predictive checks for both the training and test data (Figure 4). All of the other count models that we considered exhibited lack of fit to at least one of these statistics in posterior predictive checks. Hereafter, we report results from the zero-inflated negative binomial model.

**Table 1.** Performance of count models on the test set in descending order. Posterior means are provided with standard deviations in parentheses.

| Model | Holdout log likelihood |
| --- | --- |
| ZI Negative binomial | -3671 (70) |
| ZI Poisson | -4093 (77) |
| Negative binomial | -4298 (114) |
| Poisson | -4572 (139) |

**Figure 4.**
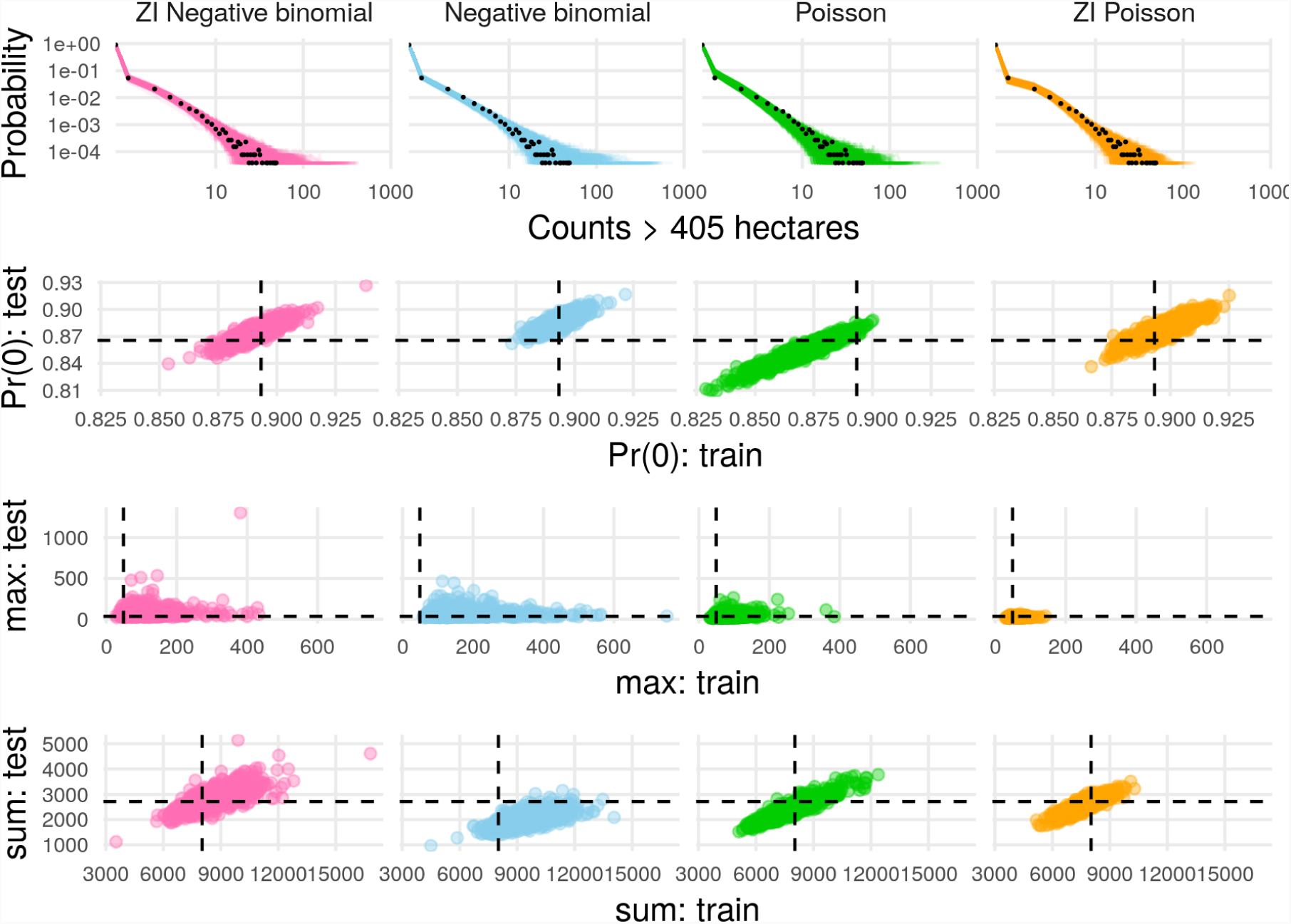
Count predictive checks. Row one shows observed count frequencies as black points and predicted frequencies as lines. Rows two, three, and four show predicted proportions of zeros, maxima, and sums (respectively) in the training and test data, with empirical values as dashed lines. Rows two through four facilitate comparison of performance on training and test sets. Ideally, model predictions cluster around the dashed lines for both the training (x-axis direction) and test (y-axis direction) sets, leading to a tight cluster of points at the intersection of the dashed lines.

Minimum relative humidity and maximum air temperature had the strongest effects on both the zero-inflation component and the expected value of the negative binomial component (Figure 5, posterior median for *ρ*: 0.665, 95% credible interval (CI): 0.319 - 0.861). The model uncovered unique effects of meteorological variables at level 1, 2, and 3 ecoregions (Figure 6). For example, a positive interaction effect between the second air temperature basis vector and the L1 Great Plains ecoregions indicates that the expected number of wildfires in plains ecoregions with cold conditions is high relative to other ecoregions. The Ozark/Ouachita-Appalachian forest and Ozark Highlands were also identified as having region-specific temperature effects (Figure 6). Twelve month total precipitation also had region specific effects in the Mississippi Alluvial and Southeast Coastal Plains ecoregion, where it was associated with lower expected fire counts (Figure 6). In contrast, increasing cumulative twelve month precipitation was associated with higher counts in desert ecoregions (Figure 5). Housing density showed a unimodal relationship to expected count (Figure 5), with lower expected counts in sparsely populated ecoregions, and higher expected counts with moderately populated ecoregions.

**Figure 5.**
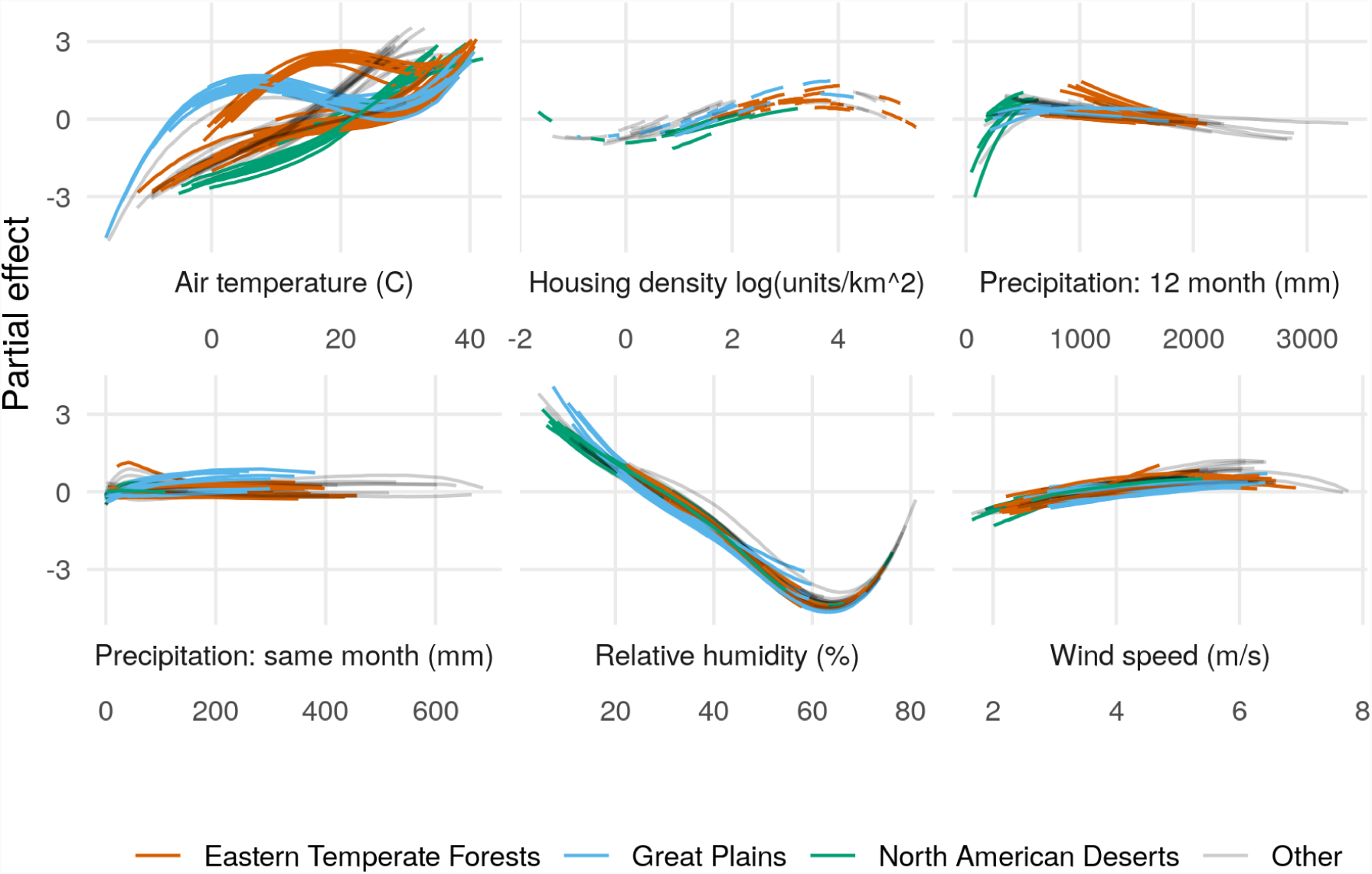
Partial effects on the log-transformed negative binomial mean component of the zero-inflated negative binomial model for each level 3 ecoregion, colored by level 1 ecoregion. Lines are posterior medians. Results are similar for the zero-inflation component.

**Figure 6.**
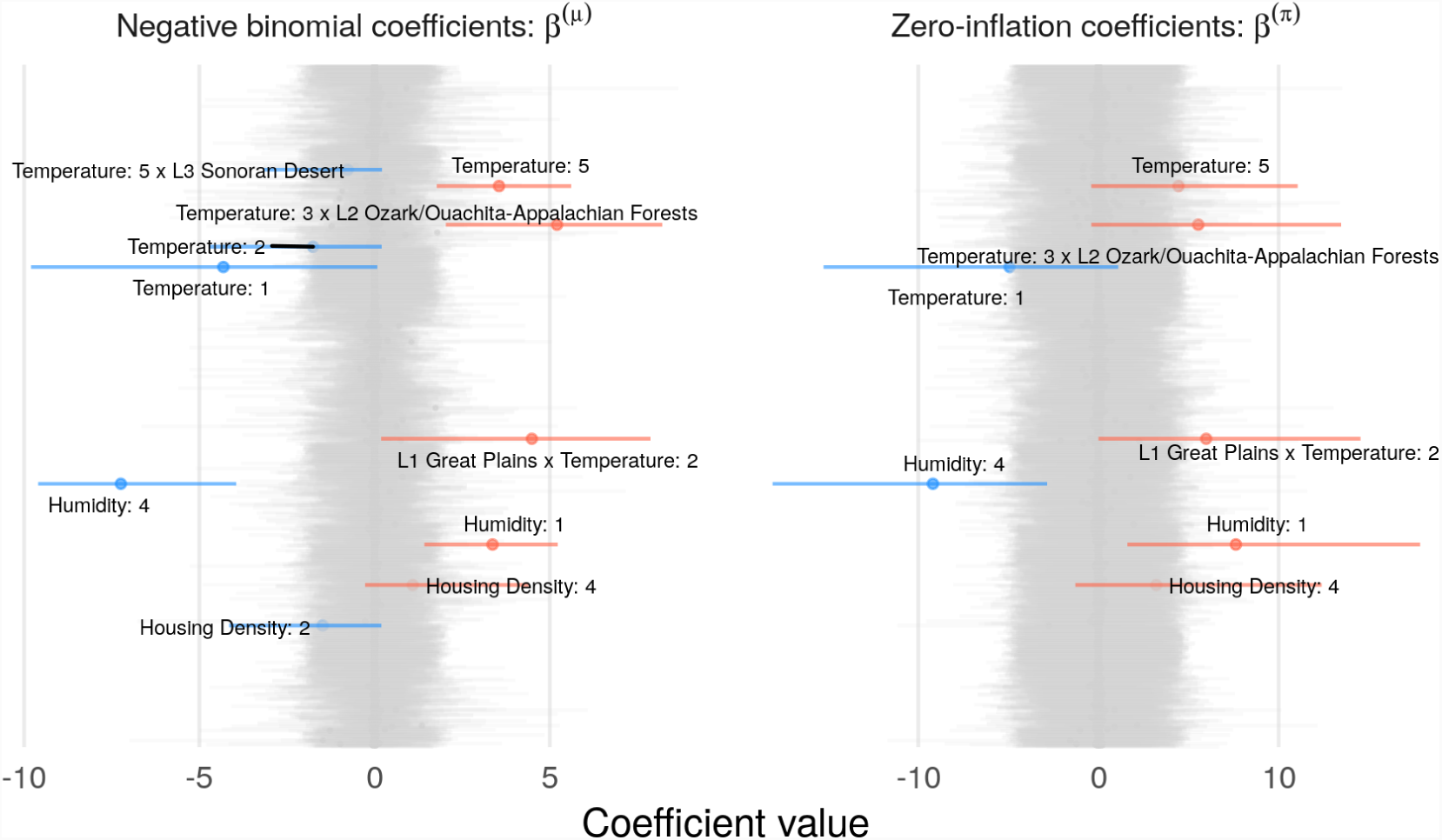
Caterpillar plots of zero inflated negative binomial model coefficients, *β*^(*µ*)^ (left) and *β*^(*π*)^ (right). Horizontal line segments denote 95% credible intervals. Grey segments indicate coefficients with a less than 87% posterior probability of being positive or negative, and colored segments indicate coefficients that are probably positive (red) or negative (blue). B-spline vectors are indicated by colons, e.g., `Humidity:1` indicates the first basis vector corresponding to humidity. Interactions between variables `a` and `b` are represented as `a x b`. Level 1 ecoregions are represented by `L1 ecoregion name`, and `L2` and `L3` indicate level 2 and 3 ecoregions.

Posterior 95% credible interval coverage for the number of fires over 405 hectares in the test set was 98.8%. The lowest test set interval coverage was 89.3%, in the Cross Timbers L3 ecoregion. When observed counts fell outside the 95% prediction interval, counts were larger than predicted 100% of the time. The largest difference between observed numbers and predicted 97.5% posterior quantiles (the upper limit for the 95% credible interval) occurred for the Columbia Mountains/Northern Rockies L3 ecoregion in August 2015, when 36 fires over 405 hectares occurred and at most 22 were predicted. For nearly half of the level 3 ecoregions (43 of 85), accounting for 39.7% of the land area of the contiguous U.S., the zero-inflated negative binomial model had 100% test set prediction interval coverage.

### Wildfire burned areas

The lognormal distribution performed best on the test set (Table 2), and captured tail-behavior better than other burned area distributions (Figure 7). The GPD model was too heavy-tailed to adequately capture the pattern in the empirical data, predicting fires far larger than those observed in the training and test sets (Figure 7). The tapered Pareto distribution was too light-tailed (Figure 7). The gamma and Weibull models performed very poorly overall on the test set (Table 2), apparently due to a lack of congruence between the shapes of these distributions and the actual burned area distribution. Despite a poor fit to the bulk of the wildfire burned area distribution, both performed adequately in the upper tails (Figure 7). Hereafter we present results for the lognormal model, which had the highest test set log likelihood and captured tail behavior of the empirical fire size distribution.

**Table 2.** Performance of burned area models on the test set in descending order. Posterior means are provided with standard deviations in parentheses.

| Model | Holdout log likelihood |
| --- | --- |
| Lognormal | -26341 (43) |
| Generalized Pareto | -26377 (45) |
| Tapered Pareto | -26386 (49) |
| Weibull | -27592 (236) |
| Gamma | -30675 (993) |

**Figure 7.**
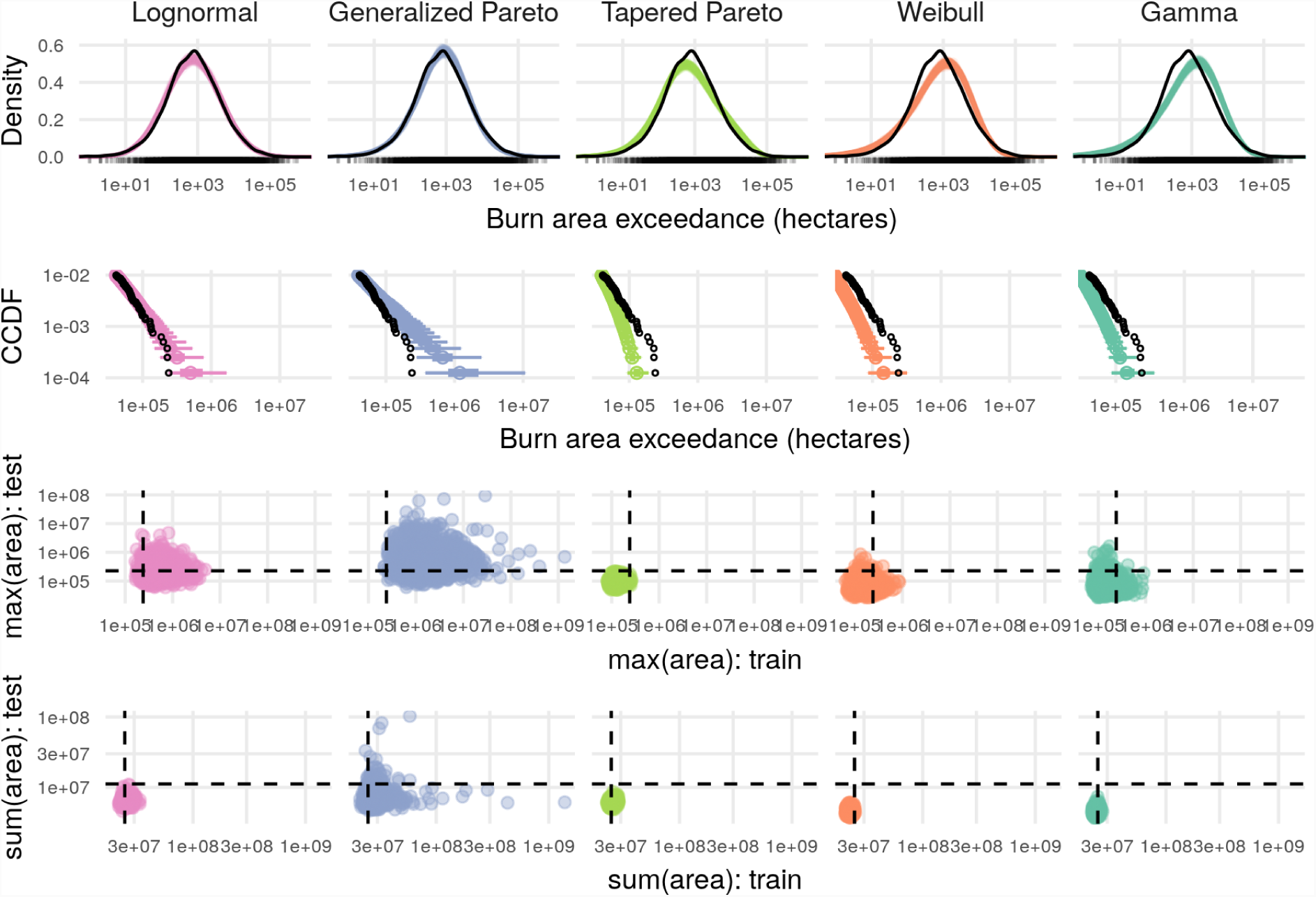
Predictive checks for burned area models. The top row shows predicted density in color and empirical density for the training set in black, which reveals overall lack of fit for the gamma and Weibull models. Row two shows the complementary cumulative distribution function (CCDF) at the tails, with 95% and 50% prediction intervals shown in color and observed data as black points, which shows that the Generalized Pareto distribution predicts values that are too extreme. The third and fourth rows show checks for maximum and total burned areas in the training and test set, with observed values as dashed lines and posterior draws as colored points. These final two rows facilitate checks for summary statistics on both the training and test set, with the ideal model generating predictions (colored points) clustered close to where the dashed lines intersect.

Relative humidity was the primary driver of expected burned area for a fire event (Figure 8A). The first basis vector for mean daily minimum relative humidity was the only coefficient with a 95% credible interval that did not include zero (posterior median: 1.68, 95% CI: (0.8 - 2.29)). This nonlinear effect can be observed in Figure 8B as an increase in the expected burned area below 20% mean daily minimum humidity. This leads to a seasonality gradient among ecoregions of expected fire sizes, with little or no seasonal signal in typically humid ecoregions such as Marine West Coast Forests of the Pacific Northwest, and seasonal oscillations in ecoregions that have periodic fluctuations between dry and humid conditions such as the Temperate Sierras (Figure 8C). There was not strong evidence that meteorological variables had spatially variable effects on expected wildfire burned area.

**Figure 8.**
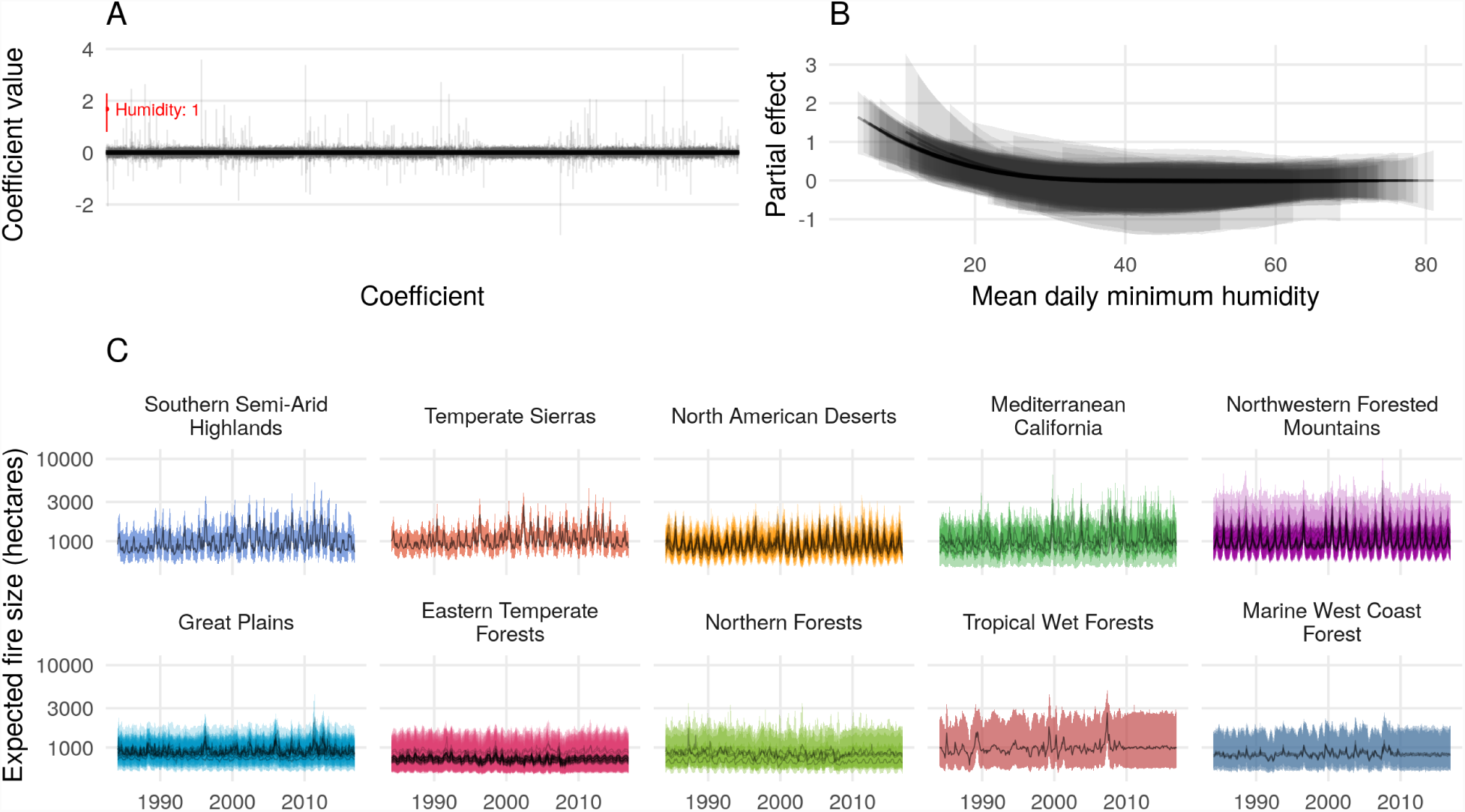
**A.** Estimated posterior medians and 95% credible intervals for each of the 3,473 coefficients associated with expected burned area. Only one coefficient the first basis vector for humidity had a 95% credible interval that excluded zero, shown in red. This effect is visualized in **B**. Partial effects of mean daily minimum humidity for each level 3 ecoregion, with posterior medians drawn as lines, and the 95% credible intervals as ribbons. **C**. Monthly time series of expected fire sizes for every level 3 ecoregion, faceted and colored by level 1 ecoregions sorted by mean humidity. Lines are posterior medians and ribbons are 95% credible intervals.

Overall, 95% posterior predictive interval coverage in the test set for burned areas was 93%. The lowest test set coverage was 0%, for the Eastern Great Lakes Lowlands L3 ecoregion, followed by 50%, for the Central California Valley L3 ecoregion, though these ecoregions had just 1 and 2 wildfire events in the test set. When observed fire sizes fell outside the 95% prediction interval, 24.9% of wildfires were smaller than predicted, and 75.1% of wildfires were larger than predicted. The largest discrepancy between the actual size of a wildfire and the predicted 97.5% posterior quantile was observed with the Wallow Fire in 2011 which burned 228,107 hectares, but the predicted upper limit for size was 20,756. We investigate this discrepancy further in the case study below. The lognormal burned area model achieved 100% interval coverage in 24 of 67 ecoregions that had wildfire events in the test set, accounting for 26% of the land area of the contiguous U.S.

### Inference on extremes

By combining the output of the event count and burned area models, we derived posterior prediction intervals for the size of the largest fire in a month for each region (the “burned area maximum”), integrating over uncertainty in the number of fires, as well as the lognormal mean and standard deviation for burned area. We evaluated the posterior distribution for the quantile function of the finite sample maximum of a lognormal distribution 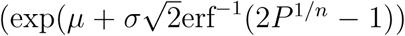, where *n* is the number of wildfire events, erf^−1^ is the inverse error function, and *P* is a probability) to generate prediction intervals for maximum fire sizes by month and ecoregion, conditional on one or more fires having occurred. In the holdout period from 2010 to 2016, a 99% prediction interval achieved 77.4% interval coverage, with 14.8% of the burned area maxima (140 fire events) being larger than predicted (Figure 9). As an additional check, we used the posterior distribution of the model to predict the total area burned by wildfires in the test set. The model predicted the total area burned over the entire contiguous United States in test period from 2010 to 2016 to be 30,339,123 (95% CI: (20,496,551 - 50,446,932) and the actual value was 30,440,173.

**Figure 9.**
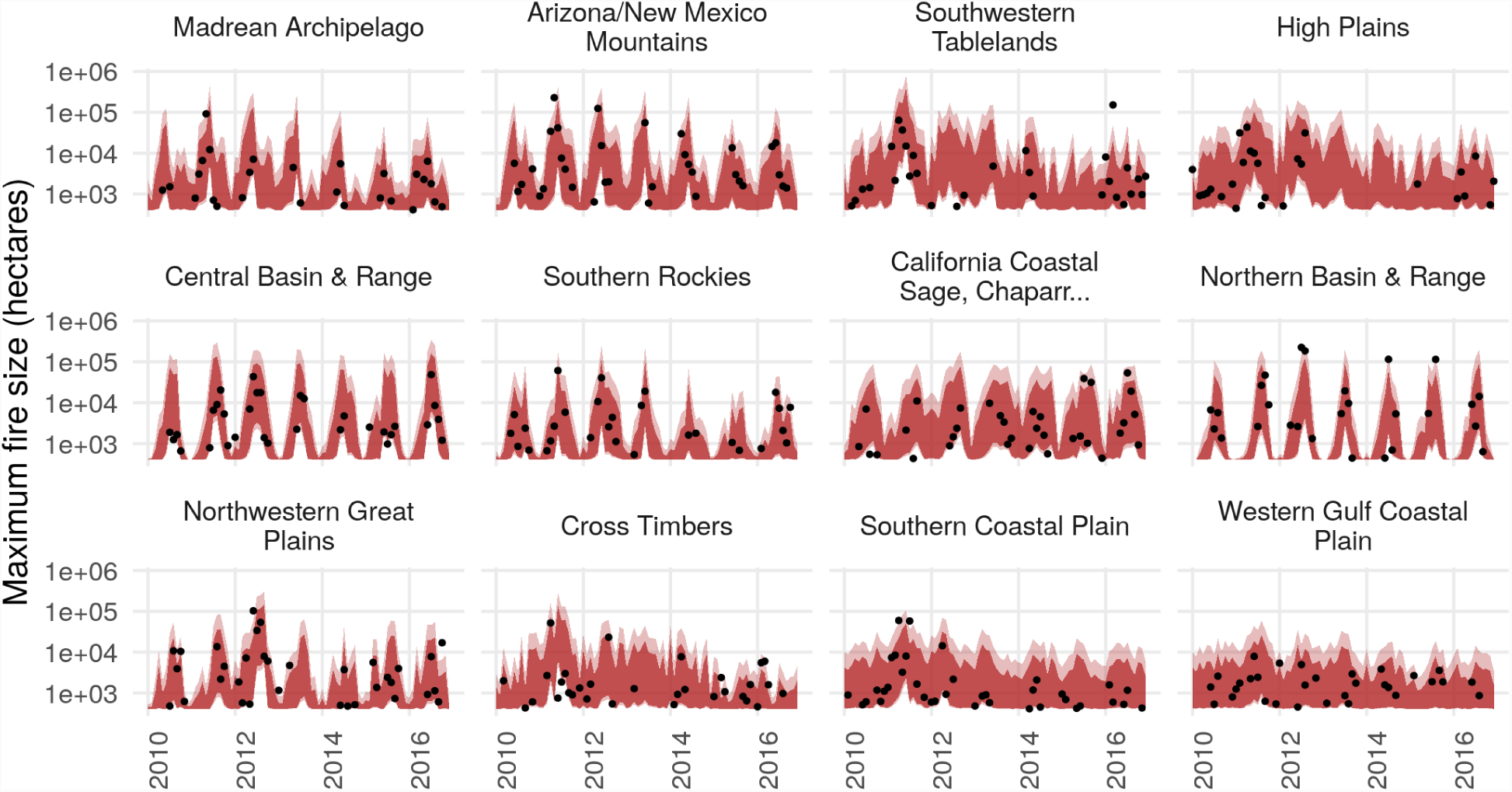
Posterior 99% (light red) and 95% (dark red) prediction intervals for the burned area of the largest fire event by month and level 3 ecoregion in the test set, shown for ecoregions with wildfires in more than 20 months. Empirical maxima are shown as black dots.

While fires over a million acres (≈ 404,686 hectares) in size have happened historically in the contiguous U.S. (Pernin 1971), no such fires were represented in in the training or test sets. If we extrapolate, the probability of at least one fire this large in the period from 2010 to 2016 was estimated to be between 0.191 and 0.651 (95% CI), with a posterior median of 0.348. The highest probability for such an event was 0.014 (posterior median), with a 95% CI of (0, 0.237) seen for the Southwestern Tablelands ecoregion in June 2011. The second highest probability was 0.004 (posterior median), with a 95% CI of (0, 0.056) seen for the Arizona/New Mexico Mountains ecoregion in June 2011. Aggregating spatially, we estimated monthly probabilities of a million acre wildfire. These probabilities show seasonal signals corresponding to peak fire seasons, with a shift toward higher and broader peaks beginning in the 21st century (Figure 10).

**Figure 10.**
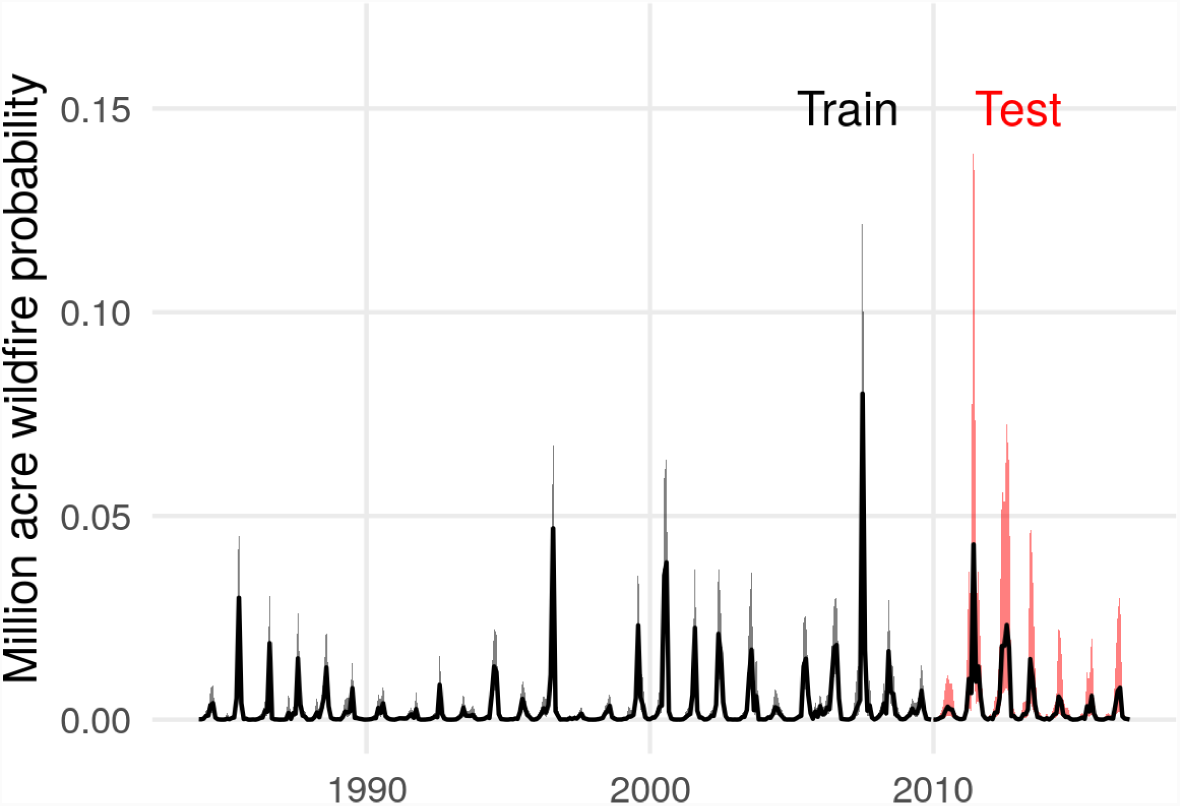
Estimated monthly posterior probabilities that one or more fire events exceed one million acres (404,686 hectares). The line represents the posterior median, and shaded region represents an 80% credible interval. The training period up to 2010 is shown in black, and the test period for which data were withheld during parameter estimation is shown in red.

### Error analysis case study: the 2011 Wallow Fire

To better understand how well the model could or could not anticipate notable extreme events, and why, we used the largest fire in the test set as a case study. The Wallow Fire was accidentally ignited on May 29, 2011 by two campers in the L3 Arizona/New Mexico Mountains ecoregion. It burned through the month of June and into early July. The model underpredicted the total burned area of the Wallow Fire. Integrating over uncertainty in the predicted number of fires and expected fire size, the 99% credible interval for the maximum fire size for May 2011 was (730 - 107,419) hectares, but the Wallow Fire is recorded as 228,103 hectares.

We evaluated the contribution of each covariate to the linear predictor functions of the three model components (lognormal mean for burned areas, negative binomial mean for counts, and the logit probability of the zero-inflation component) to understand why these predictions differed. We defined the contribution of a variable as the dot product of the elements in the design matrix **X** corresponding to a particular driver variable (e.g., humidity), and the estimated coefficients in ***β*** corresponding to that variable. This provides a quantitative measure of how each input variable contributes to the linear predictor for an ecoregion, and incorporates the overall, level 1, level 2, and level 3 ecoregion adjustments on these effects. Humidity is the primary driver of variation in the model’s predictions overall, and June 2011 the month after ignition favored more large fires, with drier, hotter conditions (Figure 11). The 99% credible interval for June 2011 was (4,258 - 428,765) hectares, which contains the true value. Had the Wallow Fire ignited two days later, the true final size would have been contained in the prediction interval. Evidently, conditions in May that drove (under)predictions of maximum burned area were not representative of the conditions over most of the Wallow Fire’s duration.

**Figure 11.**
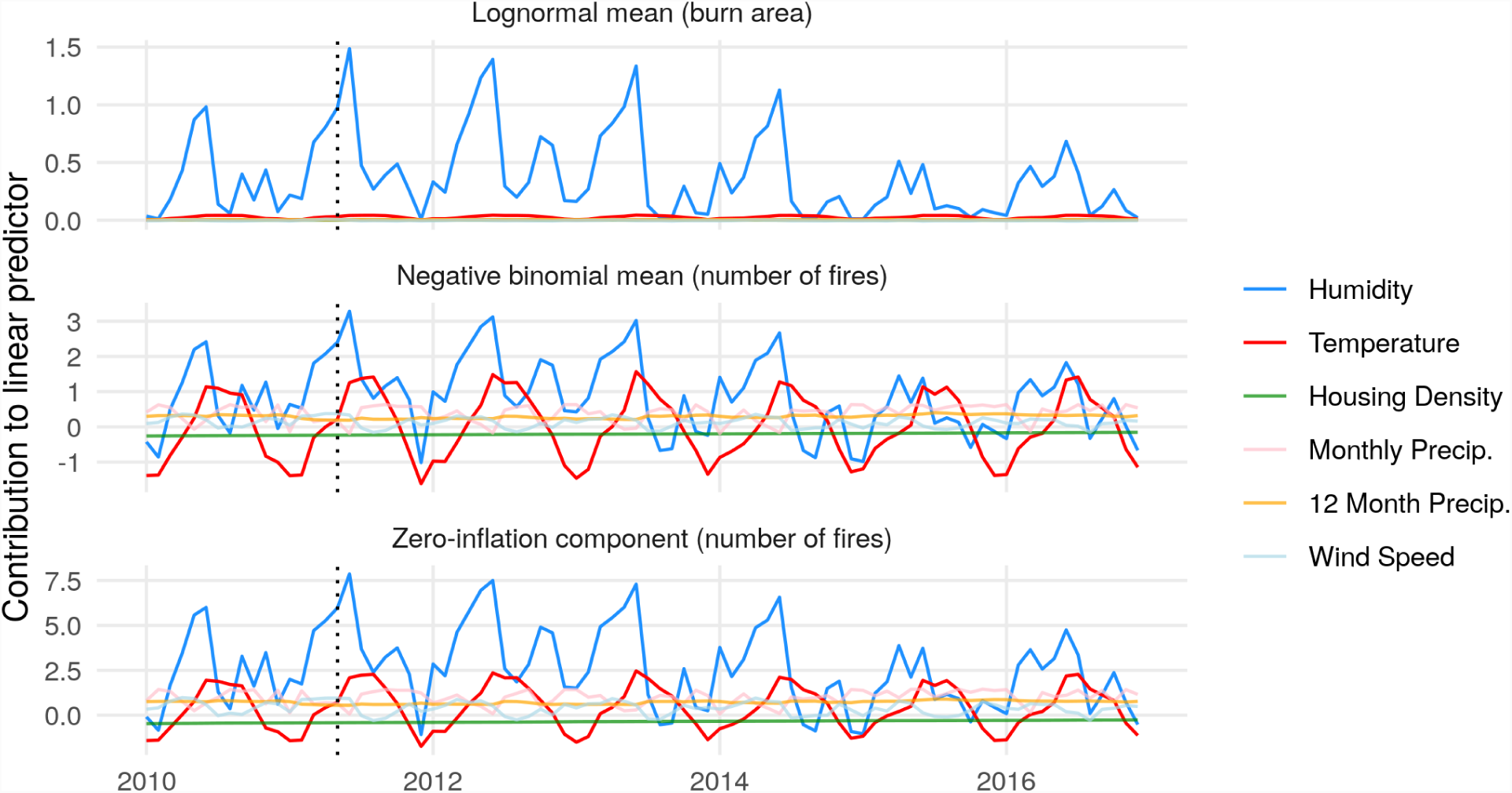
Posterior median contribution of each input variable to the linear predictor function of model components for the Arizona/New Mexico Mountains level 3 ecoregion from 2010-2016. A dotted vertical line marks May 2011, when the Wallow Fire ignited. Vertical positions of colored lines show contributions to the linear predictor function of each model component.

Temporal mismatch aside, meteorological conditions local to the Wallow Fire differed from the monthly regional means (Figure 12). In particular, wind speeds in the Wallow Fire vicinity exceeded the regional monthly mean values on the date of ignition and in the weeks following ignition. Over the majority of the duration of the Wallow Fire (May 29 to July 8), local daily conditions were drier and hotter on average than regional mean monthly conditions in May, which were used to drive the statistical model. This local variability is not represented in the regional models developed here. The failure of the model to correctly predict the size of the Wallow fire suggests potential avenues for improvement, discussed below.

**Figure 12.**
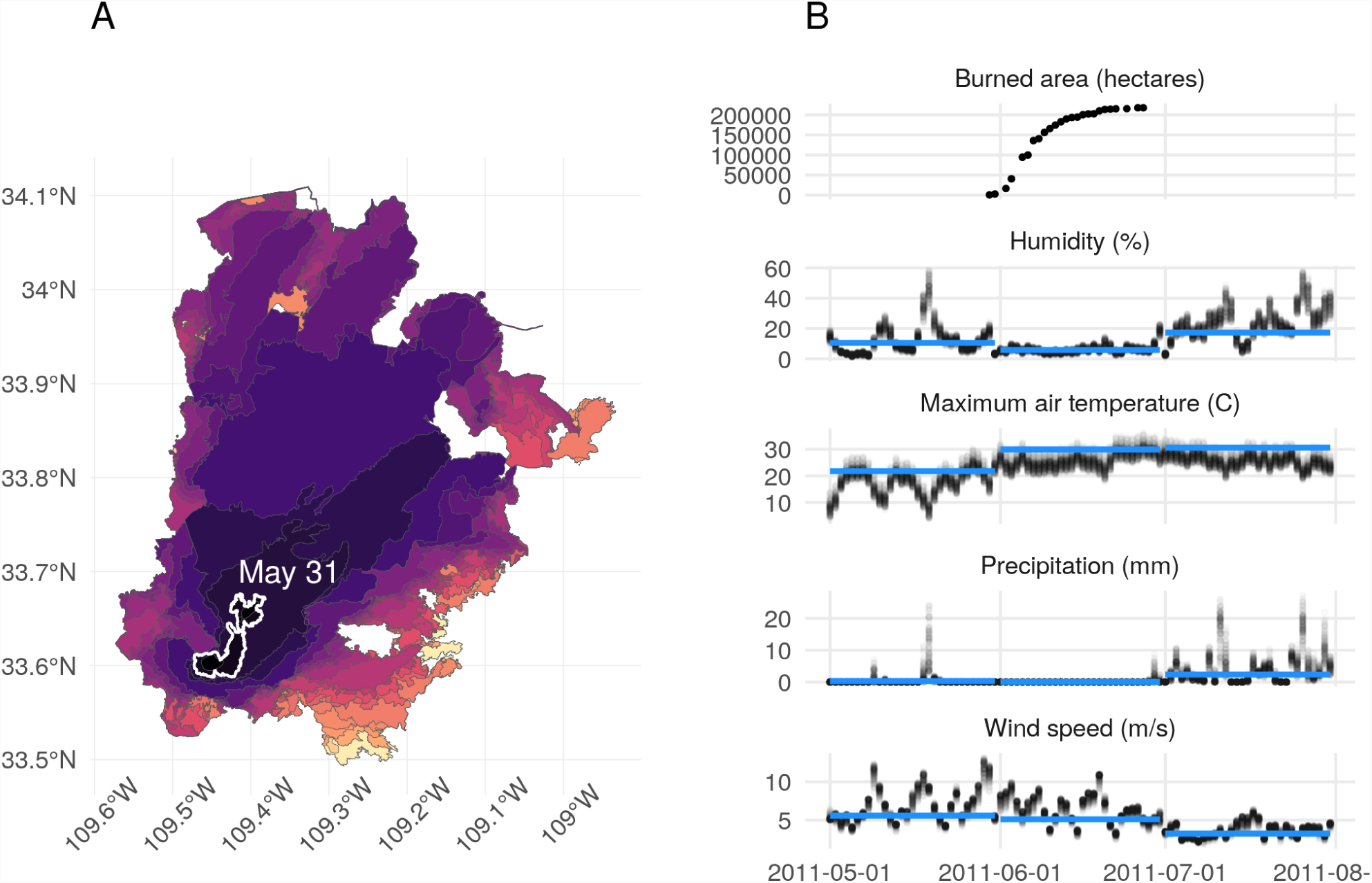
A. Progression of the Wallow Fire based on perimeter data collected from the GeoMAC database, spanning May 30, 2011 to June 27, 2011. The perimeter at the end of may is outlined in white, with brighter colors indicating later dates. B. Daily local meteorological conditions and monthly regional mean conditions for the Wallow Fire, along with the associated burned area over time. The blue line represents monthly averages of meteorological quantities computed over the entire Arizona/New Mexico Mountains ecoregion, and black points represent values extracted for “local” 4 km grid cells contained within the final burned area perimeter of the Wallow Fire.

## Discussion

Extreme wildfires are often devastating, but perhaps they need not be surprising. By allowing the non-linear effects of weather and housing density to vary across space, this model achieves good predictive accuracy for fire extremes at a regional scale over a six year prediction window. This model predicts that extremely large wildfires, perhaps even over one million acres (404,686 hectares), have a non-negligible probability of occurrence in the contiguous United States. Such predictions can support regional wildfire management and probabilistic hazard assessment.

Driving a model with meteorological features raises challenges related to predictive uncertainty and covariate shift a change in the underlying distribution of forcing variables, potentially outside of the historic range. Ideally, this uncertainty would be propagated forward in a predictive model, possibly through stacking of predictive distributions that are generated from multiple models of future climate dynamics (Yao et al. 2017). But, even if one had a perfect forecast, novel conditions present a challenge for predictive modeling (Quionero-Candela et al. 2009). For example, the High Plains ecoregion had its highest mean monthly precipitation, lowest 12 month running precipitation, driest, hottest, and windiest conditions in the test set period. Extrapolating beyond the range of training inputs is generally difficult, but the hierarchical spatial effect specification used here allows partial pooling among climatically similar ecoregions that can inform such predictions, unlike models fit separately to disjoint spatial regions.

Similar issues could arise when making predictions for observed but rare meteorological conditions. For example, mean daily minimum humidity values over 60% accounted for just 3.76% of the ecoregion-months in the training data, and 0 fires occurred in such months. As a consequence, there is relatively little data that can be used to inform the model for such conditions, and the prior distribution which shrinks coefficients toward zero may dominate the likelihood in the posterior distribution. In this case, the posterior distribution for the last basis coefficient for the partial effect of humidity is likely to be close to zero. This could explain why the estimated partial effect of humidity on the expected counts was less negative at the upper end of the observed humidity range, although previous work has found similarly nonlinear partial effects (Preisler et al. 2004). The count model performed extremely well in this range, with 100% interval coverage for the 299 ecoregion-months with mean daily minimum humidity values greater than 60% in the withheld test data. The model nearly always predicted zero counts with high confidence when conditions were this humid: 298 of 299 predictions made for such conditions were 95% credible intervals of (0, 0). The remaining prediction had a posterior median of zero, along with a 95% credible interval from 0 to 1. Monotonicity constraints could be incorporated into these models via monotonic spline bases (Ramsay and others 1988), or an ordered prior distribution for basis coefficients (Brezger and Steiner 2008). In this case, the count model performs well under humid conditions without monotonicity constraints, and there seems to be little room for performance improvements that might result from such constraints.

Human-caused climate change is expected to increase fire activity in the western U.S. (Rogers et al. 2011; Westerling et al. 2011; Moritz et al. 2012; Abatzoglou and Williams 2016) and elsewhere (Flannigan et al. 2009), but the nonlinear effect of housing density could provide additional insight into future expectations. While housing density is increasing over time in most U.S. ecoregions, some of these ecoregions are in the range of values in which this increases the expected number of large fires, while others are so populated that further increases would reduce the chance of a large fire. The hump-shaped effect of human density on the expected number of large fires is likely driven by ignition pressure and fire suppression (Balch et al. 2017). As human density increases from zero, ignition pressure increases, but eventually landscapes become so urbanized, fragmented, and/or fire-suppressed that wildfire risk decreases (Syphard et al. 2007; Bowman et al. 2011; Bistinas et al. 2013; Knorr et al. 2013; Mcwethy et al. 2013; Syphard et al. 2017; Nagy et al. 2018). At intermediate density, wildfire regimes respond to human ignition and altered fuel distributions (Guyette, Muzika, and Dey 2002), but these responses depend on environmental context and characteristics of the human population (Marlon et al. 2008; Li et al. 2009). This model indicates that the combination of moderate to high human density and dry conditions would nonlinearly increase the chance of an extreme fire event. Both human density and dryness are expected to increase in the future across large swaths of the U.S. (Lloyd, Sorichetta, and Tatem 2017; Stavros et al. 2014, Radeloff et al. (2010)), with potential implications for human mortality, health risks from smoke and particulate emission, and the financial burden of wildfire management (Reid et al. 2016; Radeloff et al. 2018).

This work points to promising directions for future predictive efforts. Default choices such as Poisson and GPD distributions should be checked against alternative distributions. Further, the predictive skill of this model seems to suggest that ordinary events provide information on extremes, which would not be the case if the generative distribution of extremes was completely unique. Previous case studies have identified that extremes or anomalies in climatological drivers play a role in the evolution of extreme wildfires (Peterson et al. 2015), but for this work, monthly averages of climatological drivers over fairly large spatial regions were used, which may smooth over anomalous or extreme conditions. Enhancing the spatiotemporal resolution of predictive models could better represent climatic and social drivers and provide localized insights to inform decision-making. This raises computational challenges, but recent advances in distributed probabilistic computing (Tran et al. 2017), efficient construction of spatiotemporal point processes (Shirota and Banerjee 2018), and compact representations of nonlinear spatial interactions (Lee and Durbán 2011) may provide solutions.

The Wallow Fire case study reveals at least one limitation of increasing the spatiotemporal resolution. When the model predictions are driven by covariates that are summarized in space and time (e.g. a mean across an ecoregion in a month), summary values may not represent conditions that are most relevant to an event. With a discrete spacetime segmentation, events can occur at the boundary of a spatiotemporal unit, e.g., if a fire spreads into an adjacent ecoregion or ignites on the last day of the month. Large wildfires can span months, and a model that only uses conditions upon ignition to predict total burned area can fail to account for conditions that change over the course of the event. Modeling ignitions as a point process in continuous space and time (Brillinger, Preisler, and Benoit 2003), and explicitly modeling subsequent fire duration and spread could better separate conditions that ignite fires from those that affect propagation. Such an approach might be amenable to including information on fuel continuity, which is likely to limit the size of extremely large fires and did not factor into the current predictions (Rollins, Morgan, and Swetnam 2002; Hargrove et al. 2000).

To the extent that a model reflects the generative process for extreme events, the decomposition of contributions to the model’s predictions may provide insight into attribution for meteorological and anthropogenic drivers of extremes. However, a model trained to represent a region-wide distribution of fire sizes will inevitably fail to capture local factors that are relevant to specific events such as the Wallow Fire. If predicting the dynamics of particular fire events is a goal, process-based models designed to model fire spread are likely to be more appropriate than regional statistical models such as those developed here.

This paper presents and evaluates a statistical approach to explain and predict extreme wildfires that incorporates spatially varying non-linear effects. The model reveals considerable differences among ecoregions spanning the mountain west to the great plains, deserts, and eastern forests, and suggests a non-negligible chance of extreme wildfires larger than those seen in over the past 30 years in the contiguous U.S. Predictive approaches such as this can inform decision-making by placing probabilistic bounds on the number of wildfires and their sizes, while provide deeper insights into wildfire ecology.

## Acknowledgments

We thank Mitzi Morris, Kyle Foreman, Daniel Simpson, Bob Carpenter, and Andrew Gelman for contributing to the implementation of an intrinsic autoregressive spatial prior in Stan. We also thank two anonymous reviewers for their thoughtful comments, which greatly improved the paper.

## Appendices

### Prior specifications

Prior distributions were chosen to regularize coefficients on the distribution specific means *β*^(*μ*)^ and structural zero parameters *β*^(*π*)^. We used a regularized horseshoe prior on these coefficients, which shrinks irrelevant coefficients towards zero, while regularizing nonzero coefficients (Piironen, Vehtari, and others 2017). For zero-inflated models, we used a multivariate version of the regularized horseshoe (Peltola et al. 2014):

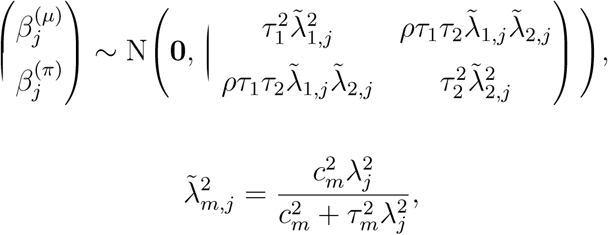

for each response dimension *m* = 1, 2 and coefficient *j* = 1, *…, p*. Here *ρ* is a correlation parameter, *τ*_1_ and *τ*_2_ are global variance hyperparameters, *c*_1_ and *c*_2_ are hyperparameters that determine the amount of shrinkage on the largest coefficients, and *λ*_*j*_ is a local scale parameter drawn from a half-Cauchy distribution that control the amount of shrinkage applied to coefficient *j* (Piironen, Vehtari, and others 2017). With this prior specification, information can be shared across the two response dimensions through the correlation parameter *ρ*, and/or through the local scale parameters *λ*_*j*_. For count models without structural zeros (the Poisson and negative binomial models), this multivariate prior simplifies to a univariate regularized horseshoe prior.

Spatiotemporal random effects were constructed using a temporally autoregressive, spatially intrinsically autoregressive formulation (Besag and Kooperberg 1995; Banerjee, Carlin, and Gelfand 2014). Temporarily suppressing the superscript that indicates whether the effects are on *μ* or *π*, and denoting column *t* from an *S* × *T* **Φ** as *ϕ*_*t*_ we have:

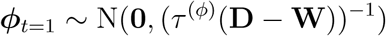

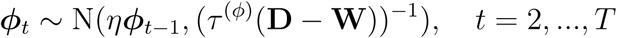

where *η* is a temporal dependence parameter, *τ* ^(*ϕ*)^ is a precision parameter, **D** is an *S* × *S* diagonal matrix with entries corresponding to the number of spatial neighbors for each spatial unit, and **W** is an *S* × *S* spatial adjacency matrix with nonzero elements only when spatial unit *i* is a neighbor of spatial unit *j* (*w*_*i, j*_ = 1 if *i* is a neighbor of *j*, and *w*_*i, j*_ = 0 otherwise, including *w*_*i, i*_ = 0 for all *i*). *τ*^(*ϕ*)^ is a precision parameter. We imposed a soft identifiability constraint that places high prior mass near 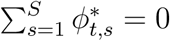 for all *t*.

We applied a univariate regularized horseshoe prior to all *β* coefficients in burned area models (Piironen, Vehtari, and others 2017):

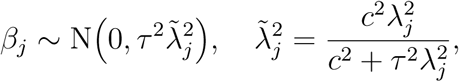

Spatiotemporal random effects were constructed in the same way as for the count models.

### Joint distributions

Here we provide the unnormalized posterior densities for each model. Square brackets represent a probability mass or density function. Parameterizations for model likelihoods are provided first, followed by the factorization of the joint distribution, with explicit priors.

#### Poisson wildfire count model

We used the following parameterization of the Poisson distribution:

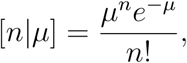

where *μ* is the mean and variance.

The unnormalized posterior density of this model is:

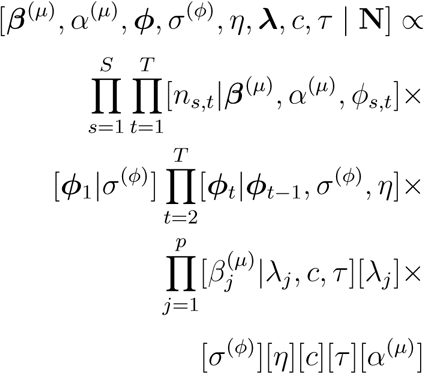

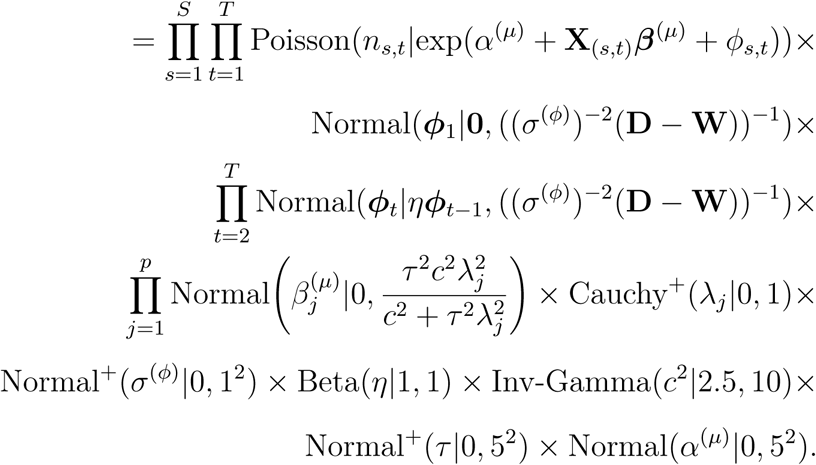

#### Negative binomial wildfire count model

We used the following parameterization of the negative binomial distribution:

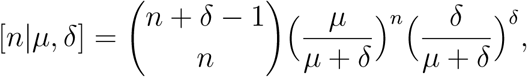

where *μ* is the mean, and *δ* is a dispersion parameter.

The unnormalized posterior density of this model is:

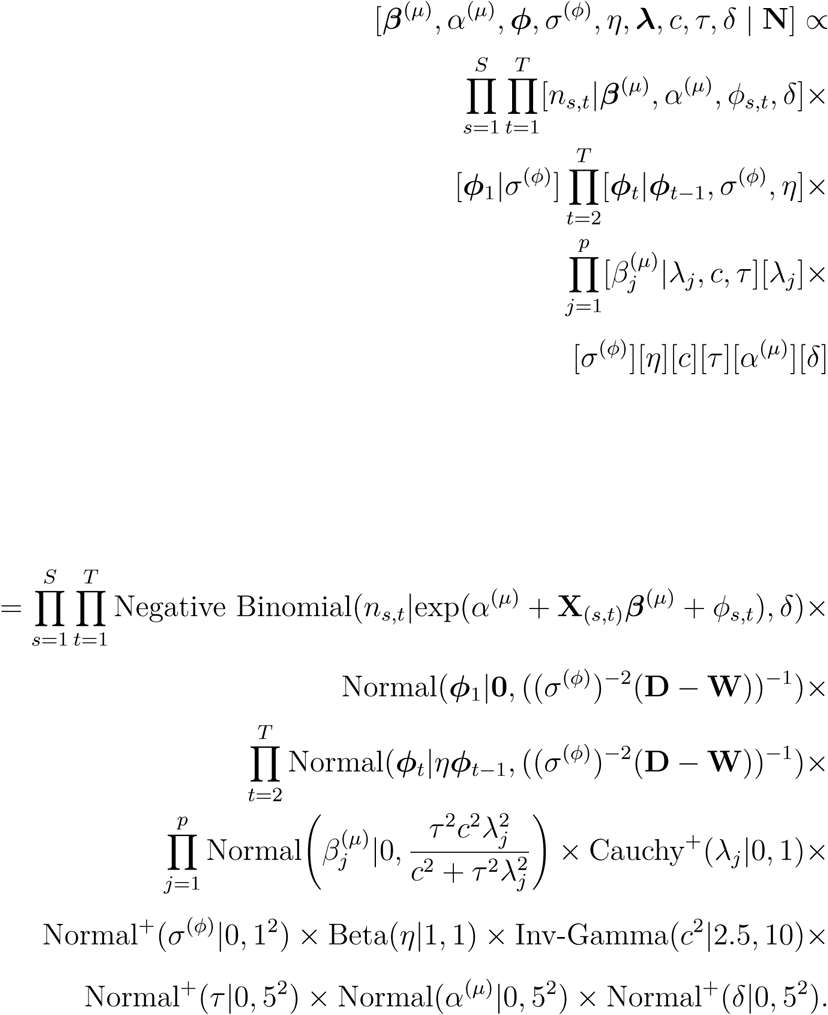

#### Zero-inflated Poisson wildfire count model

We used the following parameterization of the zero-inflated Poisson distribution:

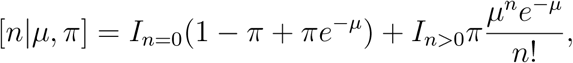

where *μ* is the Poisson mean, and 1 − *π* is the probability of an extra zero.

The unnormalized posterior density of this model is:

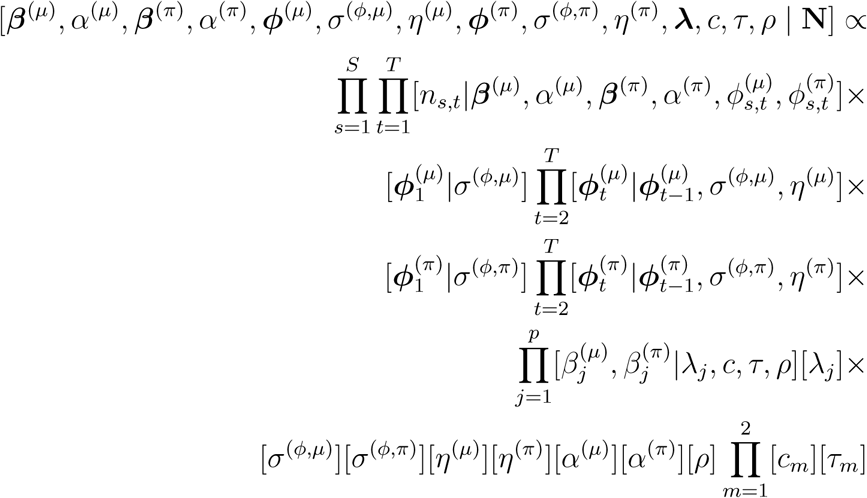

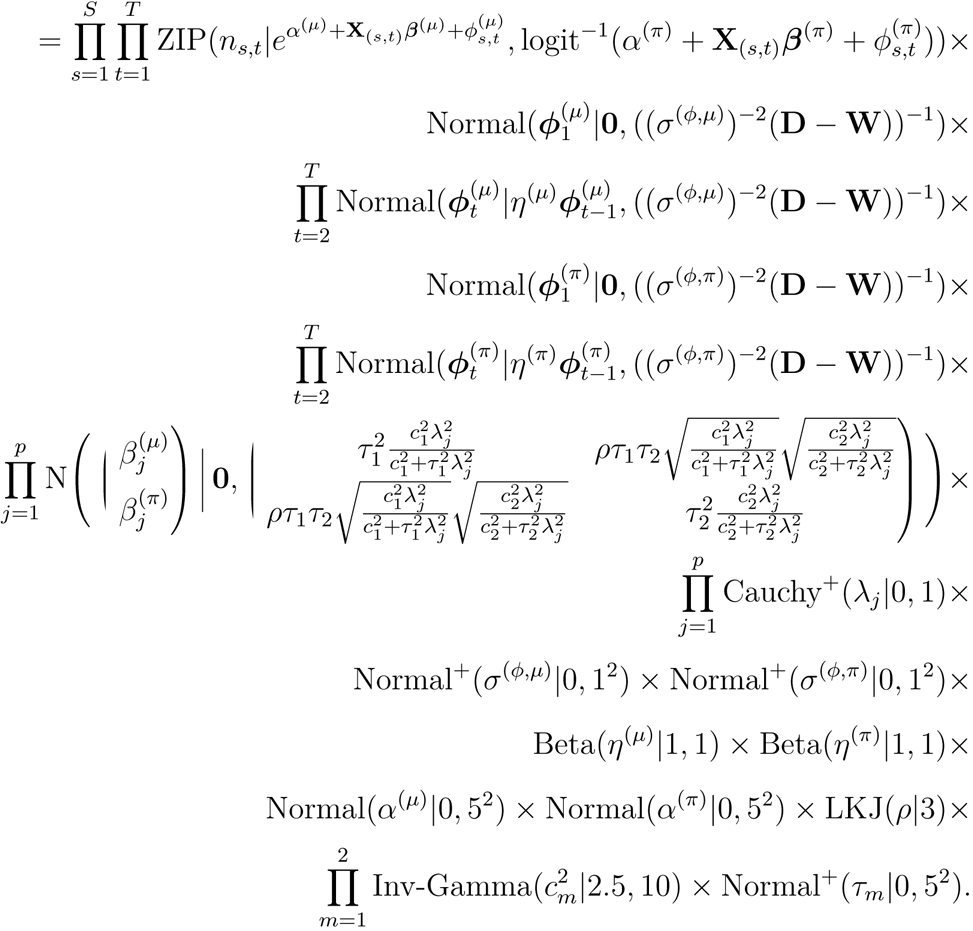

#### Zero-inflated negative binomial wildfire count model

We used the following parameterization of the zero-inflated negative binomial distribution:

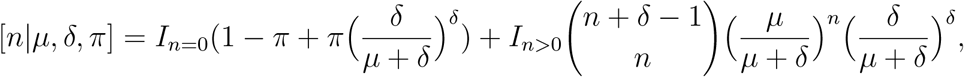

where *μ* is the negative binomial mean, *δ* is the negative binomial dispersion, and, and 1 − *π* is the probability of an extra zero.

The unnormalized posterior density of this model is:

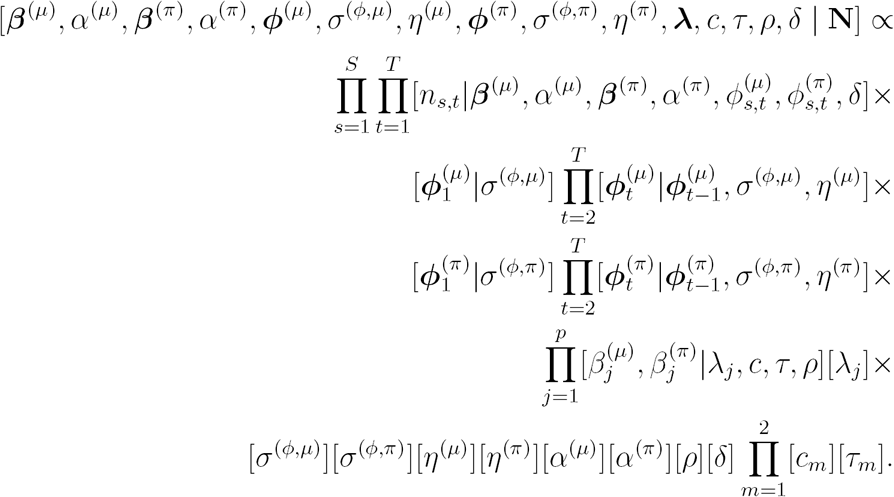

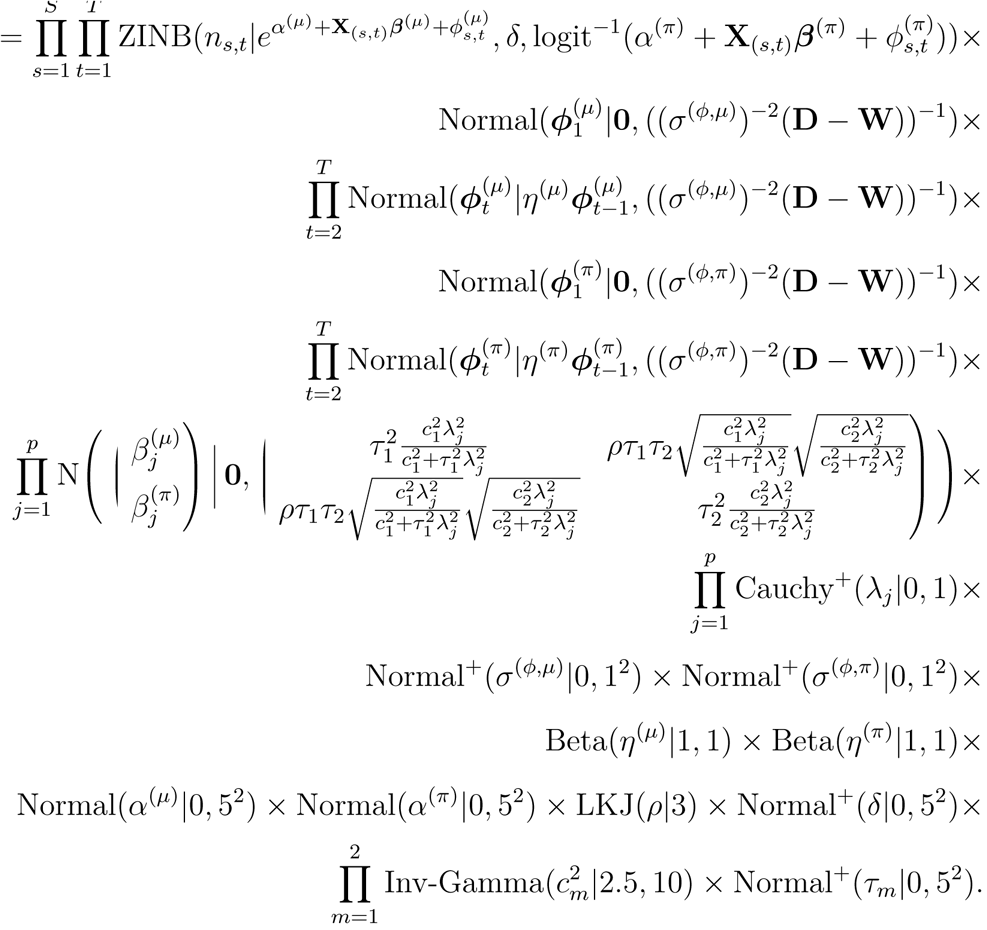

#### Generalized Pareto/Lomax burned area model

We used the following parameterization of the GPD/Lomax distribution:

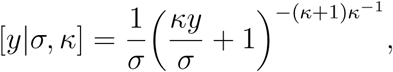

where *κ* is a shape parameter and *σ* is a scale parameter.

The unnormalized posterior density of this model is:

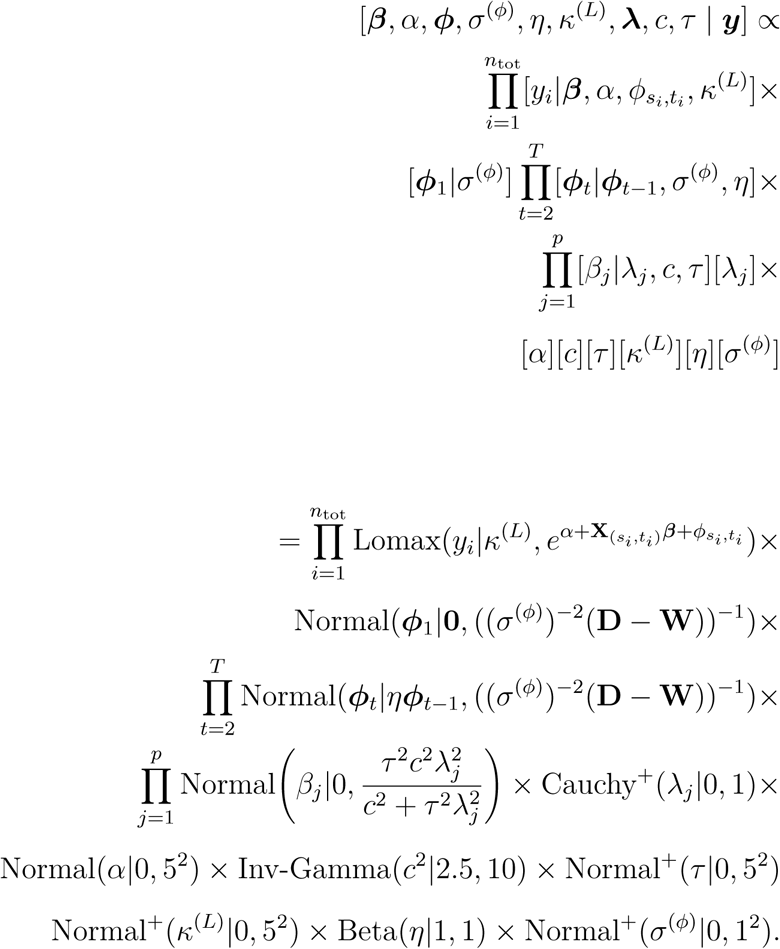

#### Tapered Pareto burned area model

We used the following parameterization of the tapered Pareto distribution:

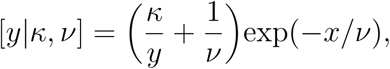

where *κ* is a shape parameter and *ν* a taper parameter.

The unnormalized posterior density of this model is:

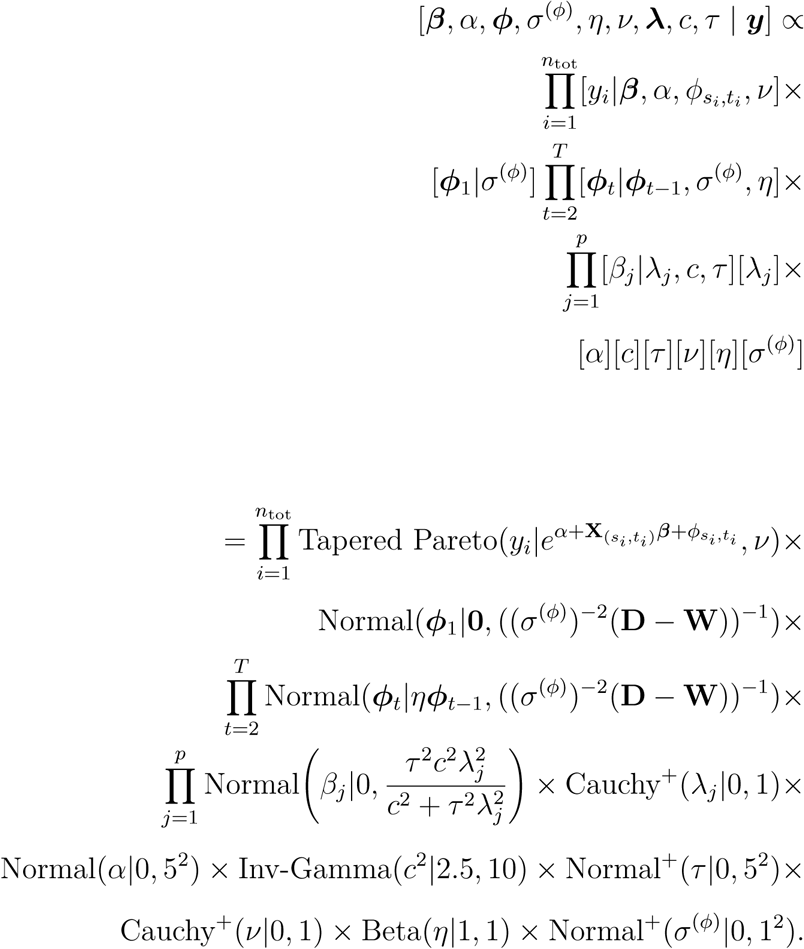

#### Lognormal burned area model

We used the following parameterization of the lognormal distribution:

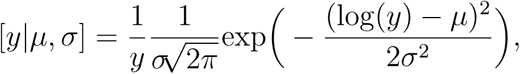

where *μ* and *σ* are location and scale parameters, respectively.

The unnormalized posterior density of this model is:

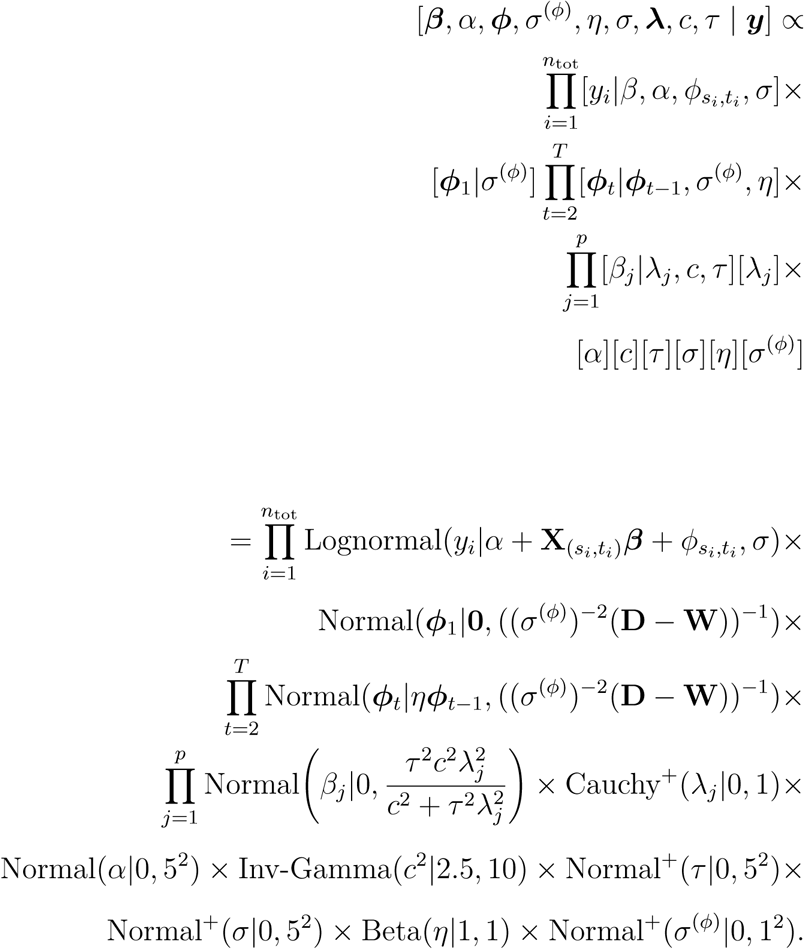

#### Gamma burned area model

We used the following parameterization of the gamma distribution:

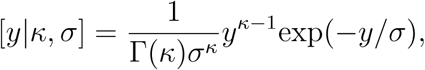

where *κ* is a shape parameter and *σ* a scale parameter.

The unnormalized posterior density of this model is:

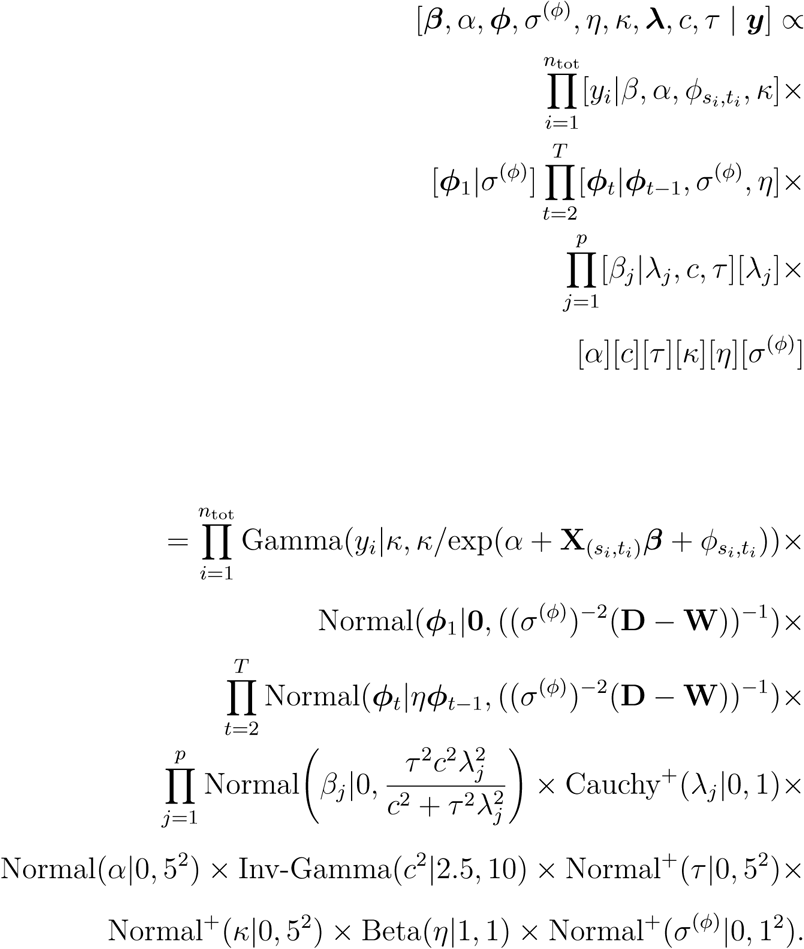

#### Weibull burned area model

We used the following parameterization of the Weibull distribution:

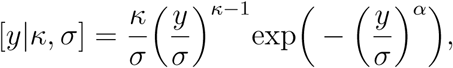

where *κ* is a shape parameter and *σ* is a scale parameter.

The unnormalized posterior density of this model is:

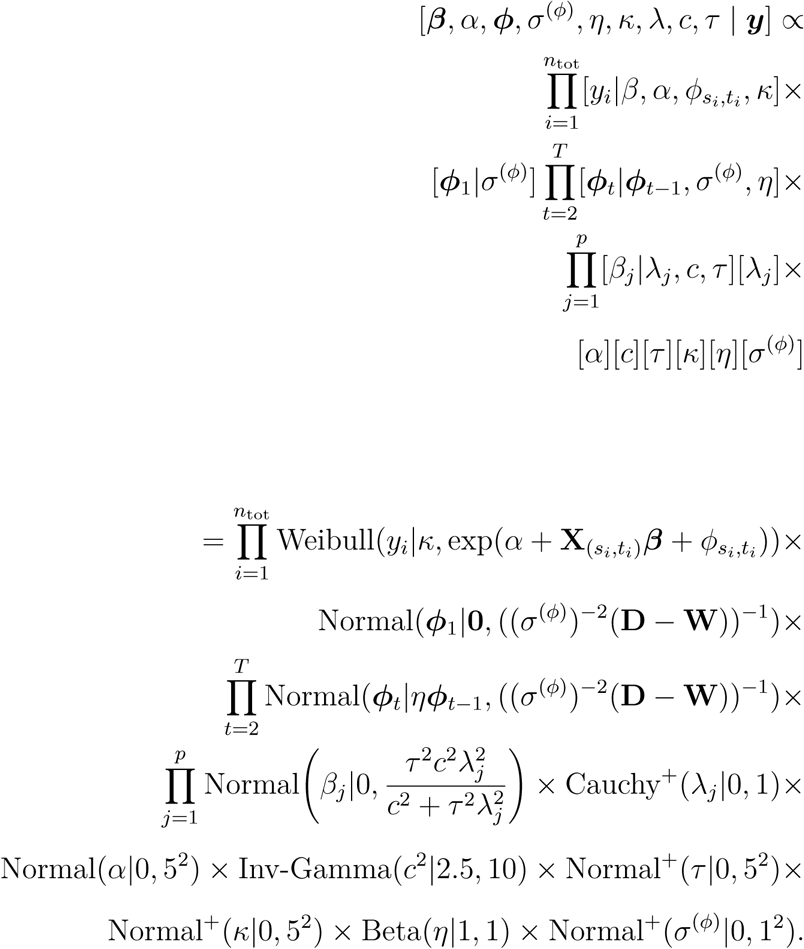

